# BRK Phosphorylates SMAD4 for proteasomal degradation and inhibits tumor suppressor FRK to control SNAIL, SLUG and metastatic potential

**DOI:** 10.1101/458190

**Authors:** Sayem Miah, Charles A. S. Banks, Yetunde Ogunbolude, Edward T. Bagu, Josh MacAusland-Berg, Anita Saraf, Gaye Hattem, Cassandra G. Kempf, Mihaela Sardiu, Scott Napper, Laurence Florens, Kiven E. Lukong, Michael P. Washburn

## Abstract

The tumor-suppressing function of SMAD4 is frequently subverted during mammary tumorigenesis, leading to cancer growth, invasion, and metastasis. A long-standing concept is that SMAD4 is not regulated by phosphorylation but ubiquitination. Interestingly, our search for signaling pathways regulated by BRK, a non-receptor protein tyrosine kinase that is up-regulated in ∼80% of invasive ductal breast tumors, led us to discover that BRK competitively binds and phosphorylates SMAD4, and regulates TGF-β/ SMAD4 signaling pathway. A constitutively active BRK (BRK-Y447F), phosphorylates SMAD4 resulting in its recognition by the ubiquitin-proteasome system, which accelerates SMAD4 degradation. In agreement, we also observed an inverse protein expression pattern of BRK and SMAD4 in a panel of breast cancer cell lines and breast tumors. Activated BRK mediated degradation of SMAD4 causes the repression of tumor suppressor genes *FRK* that was associated with increased expression of mesenchymal markers and decreased cell adhesion ability. Thus, our data suggest that combination therapies targeting activated BRK signaling may have synergized the benefits in the treatment of SMAD4 repressed cancers. Therefore, our data propose that combination therapies which includes targeting activated BRK signaling may synergize the benefits in the treatment of SMAD4 deficient cancers.

## Introduction

Breast tumor kinase (BRK) is a non-receptor tyrosine kinase highly expressed in most breast cancer cell lines and tumors(1). It displays a similar architecture and 30–40% sequence identity with Src Family Kinases (SFKs)(2). It is composed of an Src homology 3 domain (SH3 domain), an SH2 domain, and a catalytic tyrosine kinase domain(3). Like Src, BRK is negatively regulated by phosphorylation of its C-terminal tyrosine 447 and activated by phosphorylation of tyrosine 342 in the catalytic domain^4^, ^5^. Mutation of tyrosine 447 to phenylalanine significantly enhances the kinase activity of BRK(4), (5).

BRK has been implicated in several signaling cascades, notably in mitogenic signaling(6). It has been shown to enhance the mitogenic signals of EGF by promoting the activation of Akt(7). Various EGFR ligands, including EGF and heregulin, have been shown to stimulate BRK activity, resulting in increased cell proliferation and migration(7–9). Consistent with this, BRK has been shown to promote HER2-induced tumorigenesis in orthotropic transplantation-based models(10). We have also reported that BRK activation significantly enhances tumor formation in xenograft models(5). Additionally, BRK is overexpressed in over 80% of breast carcinomas(1), and in many other major cancer types including lung(11), ovarian(12), and pancreatic(13) cancers. Although the cellular role of BRK in carcinogenesis has been established, its role in controlling signal transduction pathways is still unclear.

Here, we first used a kinome array(14) to elucidate the role of BRK in regulating signal transduction pathways and identified components of the TGF-β/SMAD signaling pathway as candidate BRK targets. Although the major molecular components of the TGF-β/SMAD signaling pathway are known(15), the dynamics of TGF-β/SMAD signaling remains unclear in many systems, including normal and cancer cells. Current evidence supports that upon TGF-β/BMP receptor activation, SMAD2, and SMAD3 or SMAD1, SMAD5 and SMAD8 bind SMAD4 forming SMAD complexes which translocate into the nucleus to initiate gene regulation(16), (17). Although TGF-β/SMAD signaling networks have played pivotal roles in biological processes, disruption of their signaling has been implicated in several developmental disorders and diseases including cancers(16). The TGF-β signaling pathway can play contradictory roles during tumor development. It can function to suppress tumorigenesis, impeding the proliferation of transformed cells during the early stage of tumorigenesis(16), (18). In contrast, in some advanced cancers, loss of function mutations or low expression of SMAD2, SMAD3, and SMAD4 have been observed leading to the suppression of the TGF-β signaling pathway (16), (19) and uncontrolled cell proliferation (18). Moreover, SMAD2 and SMAD4 are being listed amongst the 127 most mutated genes in 12 major cancer types(20).

Given the complexity of TGF-β/SMAD signaling networks in biological processes, we set out to investigate how a non-receptor tyrosine kinase such as BRK may regulate the function and network of the TGF-β/SMAD signaling pathway. We used complementary immunological, biochemical, genomics, and proteomics approaches to understand how BRK might regulate TGF-β/SMAD protein interaction networks and signal transduction pathways. Collectively, our results provide evidence that BRK regulates the TGF-β/SMAD signal transduction pathway in cancer and normal cells. BRK mediated phosphorylated SMAD4 is targeted for proteasomal degradation, resulting in downregulation of the tumor suppressor Fyn-related kinase (FRK) and upregulation of the EMT markers SNAIL and SLUG in cancer cells. Taken together, our work provides a rationale for therapeutically targeting BRK in SMAD4 deficient cancer patients.

## Results

### Components of the TGF-β/SMAD pathway are potential BRK targets

BRK is overexpressed in most breast cancer cell lines and tumors(21), (5), and importantly, BRK is activated in the plasma membrane of human breast tumors(22). To substantiate the overexpression of BRK in most major cancer types, we interrogated the gene expression database, GENT (Gene Expression across Normal and Tumors; http://medicalgenome.kribb.re.kr/GENT/reference.php). We found that the expression of BRK mRNA was significantly higher (p≤ 0.05) in all five cancer types that we queried compared to their respective non-cancerous tissues (**Figure 1A**). Having confirmed that BRK overexpression is prevalent in cancers, we next sought to identify BRK targets.

**Figure 1.**
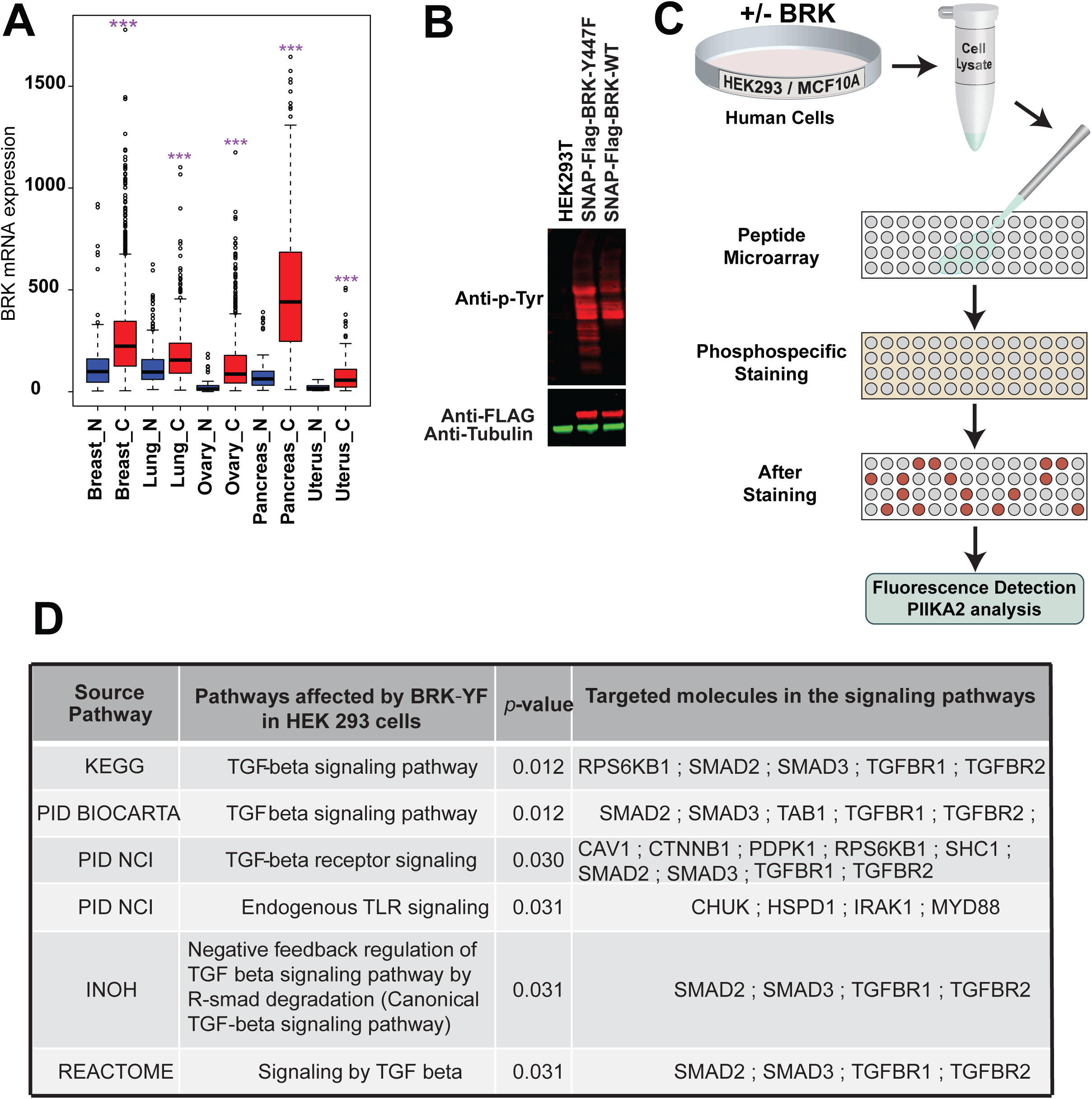
BRK is overexpressed in several human tumors and regulate different signaling pathways in normal and cancer cells. **A.** Differential expression of *BRK* in five major cancer types. Data obtained from The Cancer Genome Atlas database, median± one quartile; ^***^p <0.001. Tissue samples are denoted N for normal and C for cancer tissue. **B**. Activity of BRK-WT and BRK-Y447F (BRK-YF) mutants in transfected HEK293 cells. WT BRK and BRK-YF were transfected in HEK293 cells, and cell lysates were subjected to immunoblot with anti-phosphotyrosine antibody (PY20), and anti-BRK and anti-β-tubulin served as a loading control. **C**. Flow diagram of peptide arrays for kinome analysis. **D.** Signaling pathways significantly (p <0.05) affected by activated BRK as identified by kinome analysis in HEK293.

In this study, we focused on the constitutively-active form of BRK, BRK-Y447F (termed BRK-YF from here on). We have previously demonstrated that BRK-YF displayed higher kinase activity than BRK-WT when ectopically and stably expressed in HEK293 cells(5). To decipher the role of activated BRK in cellular signal transduction pathways, we expressed GFP-tagged, or SNAP-FLAG tagged BRK-WT and BRK-YF constructs in HEK293 cells and evaluated their global kinase activity by analyzing cell lysates by Western blotting. When we visualized phosphorylated proteins using the PY20 anti-phosphotyrosine antibody, we confirmed that BRK-YF showed higher kinase activity than BRK-WT (**Figure 1B**).

Next, we used a kinome peptide array to identify potential signaling pathways regulated by activated-BRK in BRK-YF expressing stable cells. This well-characterized kinome array(14) consisted of 300 distinct target peptides corresponding to different signaling molecules involved in various signal transduction pathways, including the TGF-β/SMAD, mTOR, PI3K, Integrin, JAK-STAT, and MAPK pathways. Lysates from stably expressing GFP-BRK-YF and parental MCF10A and HEK293 cells were analyzed using this kinome platform (**Figure 1C**). We observed that potential BRK targets were enriched for components of several signaling pathways, notably the TGFβ/SMAD signaling pathway (p ≤ 0.05), (**Figure 1D and Suppl. Fig. 1**).

### SMAD4 is a cytosolic target of BRK

As our kinome array data suggested that SMAD family proteins were potential targets for BRK mediated phosphorylation, we next asked whether SMAD2/3/4 interacted with BRK. First, we expressed GFP-SMAD2/3/4 either alone or with BRK-YF into HEK 293 cells followed by immunoprecipitation using antibodies against GFP and BRK. We found that BRK-YF co-purified with either SMAD2, SMAD3, or SMAD4 (**Figure 2A**). A reciprocal association was also observed when BRK-YF was co-expressed with either GFP-SMAD2/3/4 and immunoprecipitated with anti-BRK antibody (**Figure 2B**). Since all three of the SMAD proteins (GFP-SMAD2/3/4) interacted with BRK-YF, we next determined which of them, if any, had the strongest interaction with BRK. We co-expressed GFP-SMAD2/3/4 together with BRK-YF in HEK 293 cells and immunoprecipitated proteins from the resulting whole cell lysates with an anti-BRK antibody. We then analyzed these proteins by immunoblotting with specific antibodies against SMAD2, SMAD3, and SMAD4. We detected SMAD4 but neither SMAD2 nor SMAD3 in the BRK purified sample, suggesting that in the presence of all three SMAD proteins, SMAD4 competitively binds BRK-YF, possibly indicating a stronger affinity of SMAD4 towards BRK-YF (**Figure 2C**).

**Figure 2.**
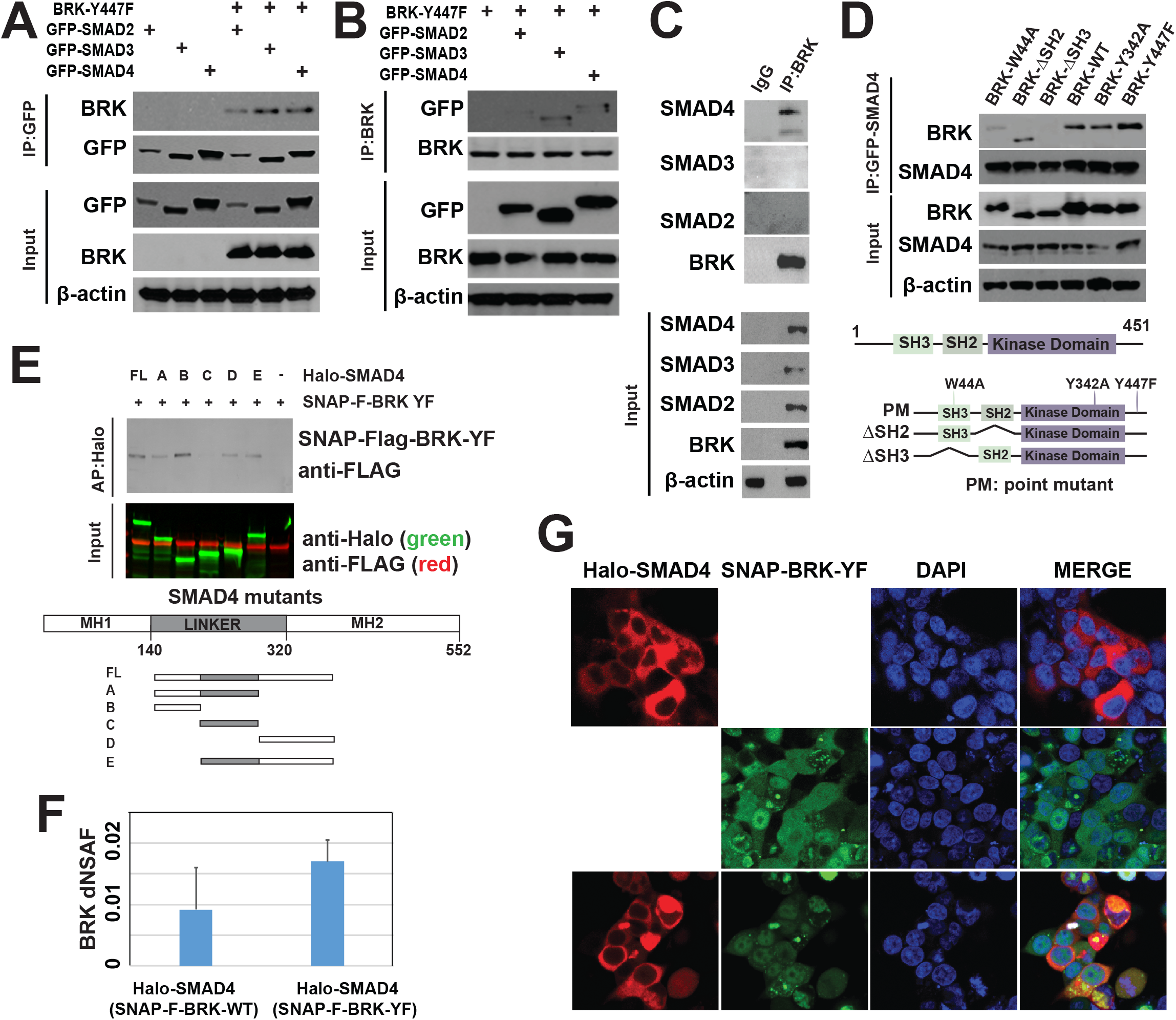
Ectopically expressed BRK and SMAD4 interact and colocalize in HEK293 cells. **A & B.** BRK-YF and GFP-SMAD2/3/4 were expressed into HEK293 cells, and cell lysates were subjected to immunoprecipitation with anti-GFP (A) or anti-BRK (B) antibodies followed by immunoblotting using anti-BRK and anti-GFP antibodies. The lower panel shows the ectopic expression of BRK and SMAD2/SMAd3/SMAD4 as detected by anti-GFP, anti-BRK antibodies; β-actin was used as a loading control. **C**. HEK293 cells were co-transfected with BRK-YF, GFP-SMAD2, GFP-SMAD3, and GFP-SMAD4 and the cell lysates were subjected to immunoprecipitation with anti-BRK followed by immunoblotting using anti-SMAD2, anti-SMAD3, anti-SMAD4, and anti-BRK antibodies. Total cell lysates were also analyzed by immunoblotting using antibodies against GFP-SMAD2, GFP-SMAD3, and GFP-SMAD4 and BRK with β-actin as loading control. **D**. GFP-SMAD4 was co-transfected with BRK-W44A, BRK-ΔSH2, BRK-ΔSH3, BRK-WT, BRK-Y342F, or BRK-Y447F. The corresponding protein extracts were immunoprecipitated with anti-GFP and immunoblotted with anti-BRK and anti-GFP antibodies and β-actin as loading control. **E**. SNAP-FLAG-BRK-YF was expressed either alone or with Halo-SMAD4 (full length: FL) or SMAD4 deletion mutants (A, B, C, D & E) in HEK293 cells. Total cell lysates were subjected to Halo affinity purification and analyzed by immunoblotting with FLAG and Halo antibodies. **F**. Halo-SMAD4 with SNAP-Flag-BRK-WT or SNAP-Flag-BRK-YF were ectopically expressed in HEK293T cells. SNAP affinity purification followed by MudPIT mass spectrometry analysis showed Halo-SMAD4 copurified with SNAP-Flag-BRK-WT or SNAP-Flag-BRK-YF. **G**. Halo-SMAD4 or SNAP-FLAG-BRK-YF alone or in combination were transfected into HEK293T cells. Halo-Tag TMRDirect fluorescent ligand (red) and SNAP-Cell^®^ 505-Star (green) were used to label Halo-Tag and SNAP-tag proteins respectively; DNA was stained with Hoechst dye (blue).

Next, to map the domains of BRK important for interaction with SMAD4, we ectopically expressed five BRK mutants with GFP-SMAD4 in HEK 293 cells. These included three mutants affecting the Src Homology domains: BRK-W44A; ΔSH2-BRK, which lacks the SH2 domain; and ΔSH3-BRK, which lacks the SH3 domain. Additionally, we used two mutants to assess whether BRK activity was necessary for interaction: the kinase-inactive BRK-Y342A and the constitutively active BRK-YF. We observed that BRK-W44A, ΔSH2-BRK, BRK-WT, BRK-Y342A, and BRK-Y447F all co-precipitated with GFP-SMAD4. In contrast, ΔSH3-BRK did not co-precipitate with GFP-SMAD4, suggesting that the SH3 domain is necessary for BRK/SMAD4 interaction (**Figure 2D**). In a similar fashion, we mapped domains of SMAD4 essential for interaction with BRK. We expressed BRK-YF in HEK293T cells together with full-length (FL) Halo-SMAD4 or with several SMAD4 truncation mutants (described in **Figure 2E**) and captured protein complexes by Halo affinity purification. We found SMAD4 mutants that contained either the MH1 domain or the MH2 domain (mutants A, B, D, and E) interacted with BRK-YF. The MH1 domain showed higher affinity for BRK-YF than the MH2 domain (compare **Figure 2E**, lanes B, and D), while the linker region alone showed a very weak affinity for BRK-YF (**Figure 2E**, lane C). Protein interactions were further analyzed by affinity purification followed by mass spectrometry (APMS) of Halo-SMAD4 ectopically expressed by itself or co-expressed with either SNAP-Flag-BRK-WT or SNAP-Flag-BRK-YF in HEK293T cells. Halo affinity purification followed by MudPIT proteomics analyses showed that Halo-SMAD4 was expressed in those cells (**Suppl. Fig. 2A**) and was able to pull down both BRK-WT and BRK-YF (**Figure 2F**). The interaction was shown to be reciprocal when SNAP-Flag-BRK-YF copurified with Halo-SMAD4 in the SNAP affinity purification analyzed by MudPIT (**Suppl. Fig. 2B**). These APMS experiments hence confirmed the conclusion from the co-immunoprecipitation experiments: BRK interacts with SMAD4.

Next, we examined whether Halo-SMAD4 and SNAP-Flag-BRK-YF co-localize *in vivo*. We used live cell imaging to assess the localization of ectopically expressed Halo-SMAD4 and SNAP-FLAG-BRK-YF in HEK 293T cells. Halo-SMAD4 and SNAP-Flag-BRK-YF were transfected in HEK 293T cells either alone or together and were imaged by confocal microscopy (**Figure 2G**). We observed that Halo-SMAD4 predominantly localized to the cytosol (**Figure 2G**, top panel), while SNAP-Flag-BRK-YF localized both in the cytosol and nucleus (**Figure 2G**, middle panel). However, Halo-SMAD4 and SNAP-FLAG-BRK-YF did colocalize in the cytosol (**Figure 2G**, bottom panel) when co-transfected in HEK 293T cells. Taken together, our observations support that SMAD4 is a cytosolic BRK interaction partner in cells, which is consistent with a potential role of SMAD4 as a target of BRK phosphorylation.

### Activated BRK phosphorylates SMAD4 on residues Tyr 353 and Tyr 412

We have shown that activated BRK interacts with SMAD4 and colocalizes with SMAD4 in the cytosol of live cells. Given that BRK is a non-receptor tyrosine kinase, we asked whether activated BRK phosphorylates SMAD4. To selectively identify BRK-mediated SMAD4 phosphorylation, we expressed Halo-SMAD4 alone or in combination with SNAP-Flag-BRK-YF in HEK 293T cell and affinity purified SMAD4 to analyze post-translational modifications (PTMs) by mass spectrometry (**Figure 3A**). Although prior studies had reported several serine, threonine, and tyrosine phosphorylation sites within SMAD4 (**Figure 3B** and not shown; source: PhosphoSitePlus^®^), our MudPIT-APMS approach identified several novel phosphorylation sites on SMAD4 in the presence or absence of SNAP-Flag-BRK-YF (**Suppl. Table 1**). Interestingly, we found that Halo-SMAD4 displayed three unique phosphorylations on S344, Y353, and Y412 in the presence of activated BRK (**Figure 3B**). Surprisingly, one of these phosphorylation sites was Serine 344, which raises the intriguing possibility that activated BRK may be a dual-specificity kinase like MEK kinases, which are involved in MAP pathways(23). Nonetheless, since BRK is a tyrosine kinase, we focused to further validate the phosphorylation of Y353 and Y412 (**Figure 3B**). We implemented a multiple reaction monitoring (MRM) approach to specifically target the peptides bearing these phosphorylated tyrosines (**Figure 3C** and **Suppl. Fig. 3A-B**). For both phosphorylated sites, the transitions of at least four fragment ions bearing the modified residues were targeted for MRM (**Figure 3C**) and were detected in the protein sample from the affinity-purified Halo-SMAD4 co-transfected with BRK-YF (**Figure 3C**). Additionally, the MS/MS spectra that were acquired immediately after the MRM spectra mapped to the expected phosphorylated peptides (**Suppl. Fig. S3A-B**).

**Figure 3.**
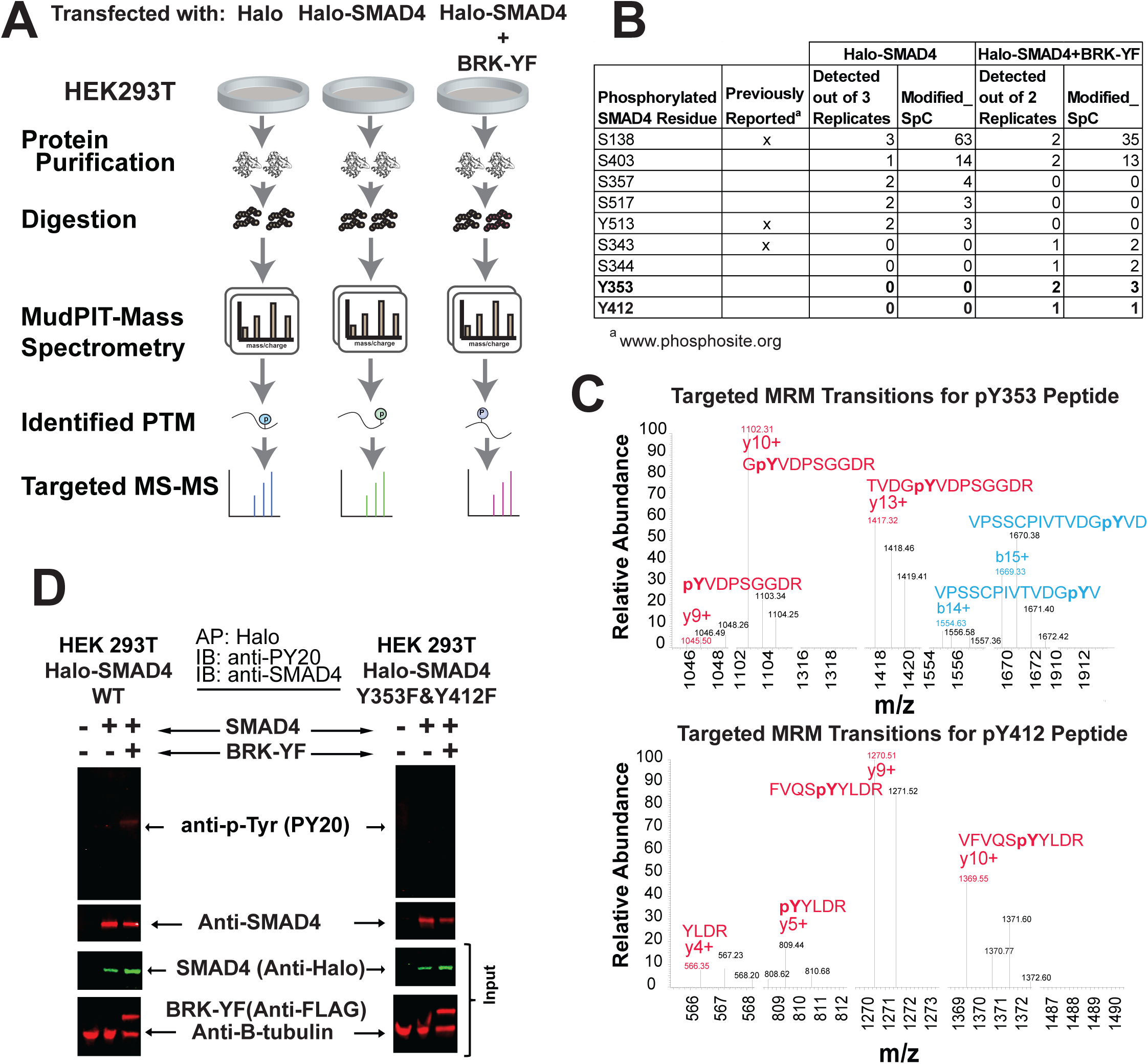
Targeted proteomics reveals BRK-mediated tyrosine phosphorylation of SMAD4. **A**. Workflow of global phosphorylation analysis by MudPIT mass spectrometry and targeted proteomics. **B**. Phosphorylation sites identified in this study (S: Serine, T: Threonine, Y: Tyrosine) tabulated with known phosphorylation sites (https://www.phosphosite.org/proteinAction.action?id=1845&showAllSites=true on Feb 7_2018). The frequency of detection and total spectral counts for the phosphorylated peptides are reported for the Halo-SMAD4 affinity purifications with or without BRK-YF. **C**. Validation of novel tyrosine phosphorylations on SMAD4 Y353 and Y412 in the presence of BRK-YF by multiple reaction monitoring (MRM). For each phosphopeptide, at least 4 fragment ions containing the modified residue were targeted for MRM. **D**. Halo-SMAD4 or Halo-SMAD4 Y353F & Y412F, with or without SNAP-Flag-BRK-YF were co-transfected into HEK293T cells and the cell lysates were subjected to affinity purification followed by immunoblotting with anti-PY20 or anti-SMAD4 antibodies. The expression of Halo-SMAD4, Halo-SMAD4 Y353F & Y412F, and SNAP-Flag-BRK-YF were analyzed by immunoblotting by using anti-Halo and anti-Flag specific antibodies. β-tubulin was used as a loading control.

To further validate BRK-mediated phosphorylation of SMAD4 Y353 and Y412, we next generated a mutant of SMAD4 lacking these two tyrosine residues (Halo-SMAD4 Y353F & Y412F). We first affinity purified Halo-SMAD4 from cells co-expressing BRK-YF with Halo-SMAD4 and analysed the pulled-down proteins with an antibody specific to phosphorylated tyrosines, PY20 (**Figure 3D**, left panel). We detected phosphorylated SMAD4 in the presence of BRK-YF, but not in the absence of BRK-YF. Repeating this experiment on the mutant Halo-SMAD4 Y353F & Y412F protein, we could not detect a similar band indicating phosphorylated SMAD4 (**Figure 3D**, right panel). Our mass spectrometry analysis of phosphorylated peptides from SMAD4 co-expressed with activated BRK-YF hence revealed two novel tyrosine residues phosphorylated explicitly by BRK. These sites were confirmed as phosphorylated with an orthogonal MRM mass spectrometry approach and by site-directed mutagenesis combined with Western blotting. These two tyrosines are located within SMAD4 MH2 domain, which is the most frequently mutated in cancers, and are likely involved in regulating the SMAD4 protein interaction network.

### BRK and SMAD4 protein expression levels are inversely correlated in most breast cancer cells and tissues

Since SMAD4 is a tumor suppressor(18), (16) and we and others have previously shown that BRK acts as an oncogene(5), (13), (9), we opted to explore the possible connection between SMAD4 and BRK protein levels in breast cancer cells and tissues. First, we examined the endogenous protein expression of SMAD4 and BRK to determine the expression profiles of these proteins in breast cancer cells. Using antibodies against each protein, we evaluated the expression level of SMAD4 and BRK in a panel of 10 breast cancer cells, two immortalized cell lines commonly used to model non-diseased human mammary epithelial cells (MCF-10A and MCF-12F), and lastly in HEK 293 cells. We detected SMAD4 in MCF-10A, MCF-12F, and HEK293, as well as in 6 breast cancer cell lines MDA231, Hs578T, T47D, BT549, MCF7, BT474. SMAD4 expression was very low or undetectable in four other breast cancer cell lines: BT20, HCC1428, SKBR3, and HCC1954 (**Figure 4A**). Detectable amounts of BRK were observed in BT20, HCC1428, SKBR3, T47D, MCF7, BT474, and HCC1954, but not in MDA231, Hs578T, BT549, and HEK 293 cells, while very low expression was observed in MCF10A and MCF12F (**Figure 4A**). Interestingly, the levels of SMAD4 and BRK were inversely correlated in MCF-10A, MCF-12F, MDA MB 231, BT20, HCC1428, SKBR3, Hs578T, BT549, HCC1954, and HEK 293. In particular, the expression of SMAD4 and BRK in all of the triple negative breast cancer (TNBC) cells we tested (MDA231, BT20, Hs578T, BT549, and HCC1954) showed this inverse expression pattern (**Figure 4A**).

**Figure 4.**
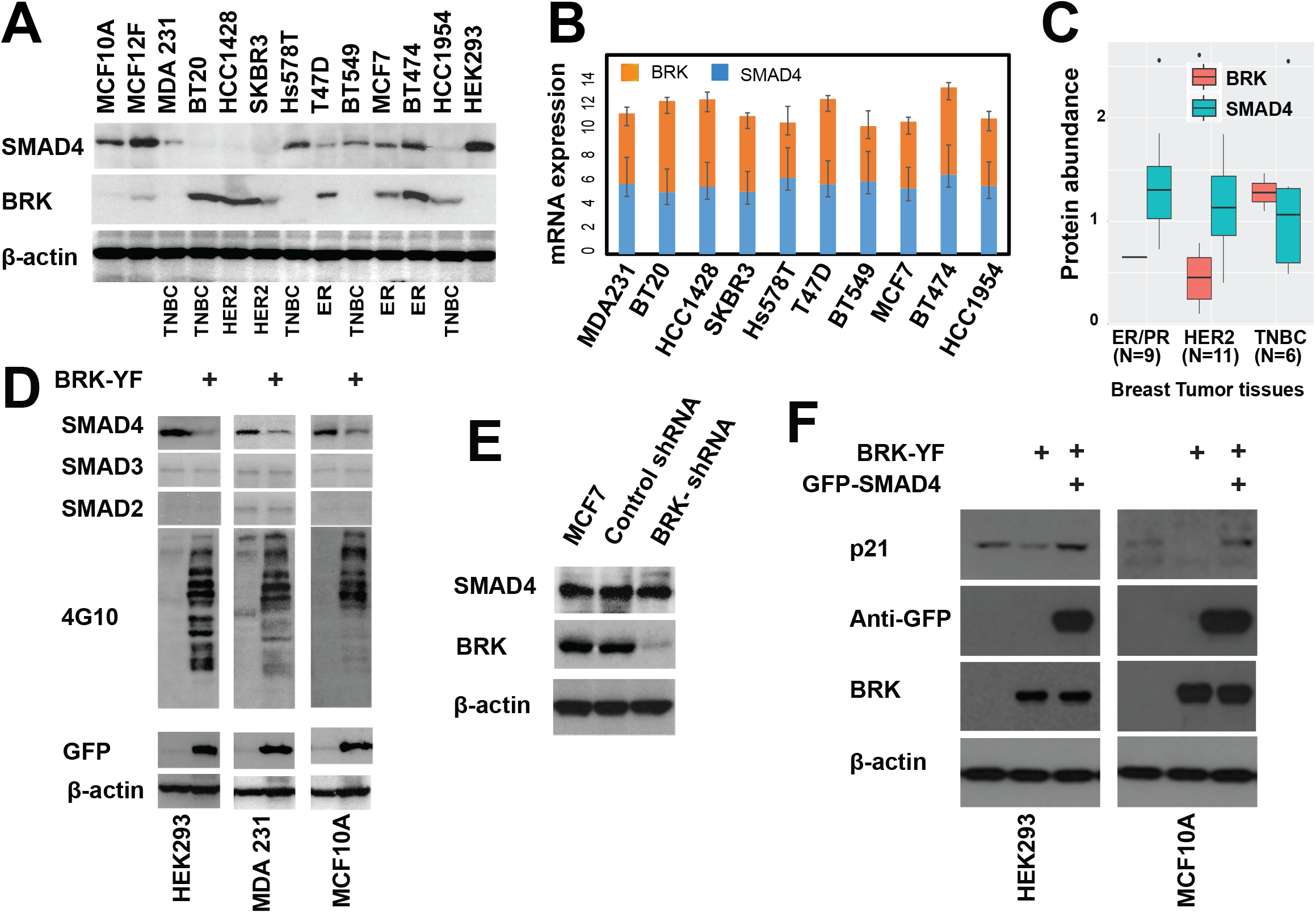
BRK and SMAD4 are expressed in most breast cancer cells and tissues. **A**. BRK expression was detected by immunoblotting in the indicated normal mammary epithelial and breast cancer cell lines (TNBC: triple negative breast cancer cell; HER2: human epidermal growth factor 1; ER: estrogen receptor), and β-Actin was used as a loading control. **B.** Differential expression of *SMAD4* and *BRK* in breast cancer cell lines, as obtained from The Cancer Cell Line Encyclopedia (CCLE). **C**. Absolute expression of BRK and SMAD4 in patients driven tumor tissues of three major breast cancer subtypes: Estrogen/Progesterone (ER/PR), HER2 and TNBC. Super-SILAC based absolute proteins expression data were obtained from Tyanova *et al.* (24). **D**. Immunoblotting analysis of total cell lysates from HEK293, MDA 231, and MCF10A cells with or without ectopically expressed GFP-BRK-YF. Stable cell lysates were analyzed by immunoblotting using SMAD2, SMAD3, SMAD4, anti-phosphotyrosine (4G10), and GFP antibodies. β-actin served as a loading control. **E**. Stable BRK knockdown was executed by using shRNA lentiviral vector plasmids (Santa Cruz Biotechnology) on the MCF7 parental breast cancer cell lines, according to the manufacturer′s protocol. Immunoblotting analysis of total cell lysates showed the expression of SMAD4 and BRK with β-actin as a loading control. **F**. Total cell lysates from HEK293 and MCF10A; GFP-BRK-YF expressing HEK293 and MCF10A stable cell lines and GFP-SMAD4 transfected GFP-BRK-YF expressing stable HEK-293 and MCF10A stable cell lines were analyzed by immunoblotting by using p21, SMAD4, and BRK specific antibodies. β-actin was a loading control.

These diverse patterns of BRK/SMAD4 protein levels might be explained by differences in BRK or SMAD4 mRNA levels. To compare the gene expression pattern of *SMAD4* and *BRK*, we examined RNAseq data from The Cancer Cell Line Encyclopedia (CCLE, https://portals.broadinstitute.org/ccle) corresponding to the 10 breast cancer cell lines that we had analysed. Surprisingly, we observed that *SMAD4* and *BRK* showed similar patterns of mRNA expression in all 10 breast cancer lines including TNBC cells (**Figure 4B**), suggesting that the effects we had initially noticed were likely regulated at the protein level rather than mRNA expression. Next, to reinforce the evidence obtained using cancer cell lines, we mined data from patient tumor samples(24) to determine SMAD4 and BRK protein expression patterns in three major breast cancer subtypes: Estrogen/Progesterone (ER/PR) positive, Human epidermal growth factor receptor 2 (HER2) positive, and triple negative breast cancer (TNBC). Again, we observed that SMAD4 and BRK were inversely expressed in these breast cancer subtypes (**Figure 4C**).

To further interrogate the relationship between SMAD4 and BRK expression, we stably-expressed constitutively-active BRK (BRK-YF) into three cell lines that expressed SMAD4 but not BRK (HEK 293, MDA-MB 231 and MCF10A cell lines). An elevated level of phosphorylation of cellular targets was observed in the cells stably expressing GFP-BRK-YF, as visualized by immunoblotting with an anti-phosphotyrosine antibody (4G10) (**Figure 4D**). We, therefore, examined the expression of SMAD4 in BRK-YF expressing cells and observed a sharp reduction of endogenous SMAD4 protein in all three of the cell lines expressing activated BRK compared to those parental cells (**Figure 4D**). However, we did not observe any noticeable change in the endogenous protein levels of SMAD2 and SMAD3 (**Figure 4D**), suggesting that the effect that activated BRK had on protein levels was specific to SMAD4.

Finally, since our data showed that SMAD4 and BRK protein levels were inversely correlated, we asked whether knocking down BRK by short hairpin RNA (shRNA) could modulate the levels of SMAD4 in MCF7 cells. MCF7 cells were selected because both SMAD4 and BRK proteins were moderately expressed in these cells (**Figure 4A**). We attained a 70–80% reduction of BRK in MCF7 cells (**Figure 4E**). However, we did not observe any noticeable effect on the expression of SMAD4 protein in MCF7 cells with depleted BRK (**Figure 4E**). Since the depletion of BRK did not affect the expression of SMAD4, we aimed to assess the expression of a known target of SMAD4 in GFP-BRK-YF cells. As a proof of principle, we evaluated the expression of p21, a known target of SMAD4(15), (25), in the cells stably expressing GFP-BRK-YF and the parental cell lines. We found that p21 protein levels sharply decreased in the cells expressing activated BRK compared to parental cells (**Figure 4F**). Moreover, reduction of p21 induced by constitutively active BRK-YF could be rescued by ectopically overexpressed GFP-SMAD4 (**Figure 4F**), suggesting that large amounts of the GFP-SMAD4 compensated for the degradation of endogenous SMAD4 mediated by BRK-YF phosphorylation, hence restoring the downstream expression of the p21 protein.

Overall, our data indicate that, compared with control cells, the increased levels of BRK observed in several breast cancer cell types was concomitant with reduced levels of SMAD4, and that changes in protein level did not necessarily result from changes in BRK/SMAD4 mRNA levels. Additionally, introducing the active BRK-YF into cells expressing SMAD4 resulted in reduced levels of SMAD4. As BRK and SMAD4 showed similar mRNA expression patterns but differ in protein levels, it is possible that BRK suppresses SMAD4 through the ubiquitin/proteasome pathway.

### SMAD4 is a target of ubiquitin modification enzymes in presence of activated-BRK

Since we had found that BRK phosphorylates SMAD4 (**Figure 3**) and SMAD4 levels were dramatically lower in stably expressing BRK-YF cells (**Figure 4D**), we next examined whether the presence of BRK-WT/BRK-YF made Halo-SMAD4 a potential target of the ubiquitin-proteasome machinery. To interrogate the impact of constitutively active BRK on SMAD4 ubiquitination, we used our established workflow(26) for Halo/MudPIT APMS analysis (**Figure 5A**). We first expressed Halo-SMAD4 in the presence or absence of SNAP-Flag-BRK-WT or SNAP-Flag-BRK-YF and purified Halo-SMAD4 associated proteins by affinity chromatography. We identified SMAD4 associated proteins by MudPIT mass spectrometry (**Suppl. Fig. S4A-C**). Indeed, we found that Halo-SMAD4 recruited several ubiquitin and deubiquitin ligases in the presence of BRK-WT/ BRK-YF (**Figure 5B and Suppl. Table 2**). SMAD4 interacted with the deubiquitin ligases USP9X, USP32, and USP7 in the presence of both BRK-WT and BRK-YF. However, SMAD4 association with the ubiquitin ligases HERC2 and RNF138 were upregulated only in the presence of BRK-YF. Since Halo-SMAD4 interacted with ubiquitin ligases in the presence of activated BRK, we then tested the possibility that the BRK-mediated phosphorylation of SMAD4 facilitated its degradation through the ubiquitin/proteasome system(27). Halo-SMAD4, BRK-YF, and HA-ubiquitin were expressed in HEK 293T cells in the combinations shown in **Figure 5C** and cells were treated with or without the peptide-aldehyde proteasome inhibitor MG132 (carbobenzoxyl-L-leucyl-L-leucyl-L-leucine). We examined SMAD4 protein for ubiquitination by resolving Halo affinity purified samples by SDS-PAGE and immunoblotting the resulting Western blot using anti-HA antibody. Our data showed a smear of ubiquitin-conjugated SMAD4 in the presence of proteasome inhibitor (**Figure 5C**, lane 5), which was not apparent in controls (**Figure 5C**, lanes 1–4). Our results led us to conclude that the presence of activated BRK causes the downregulation of SMAD4 by the ubiquitin/proteasome degradation pathway.

**Figure 5.**
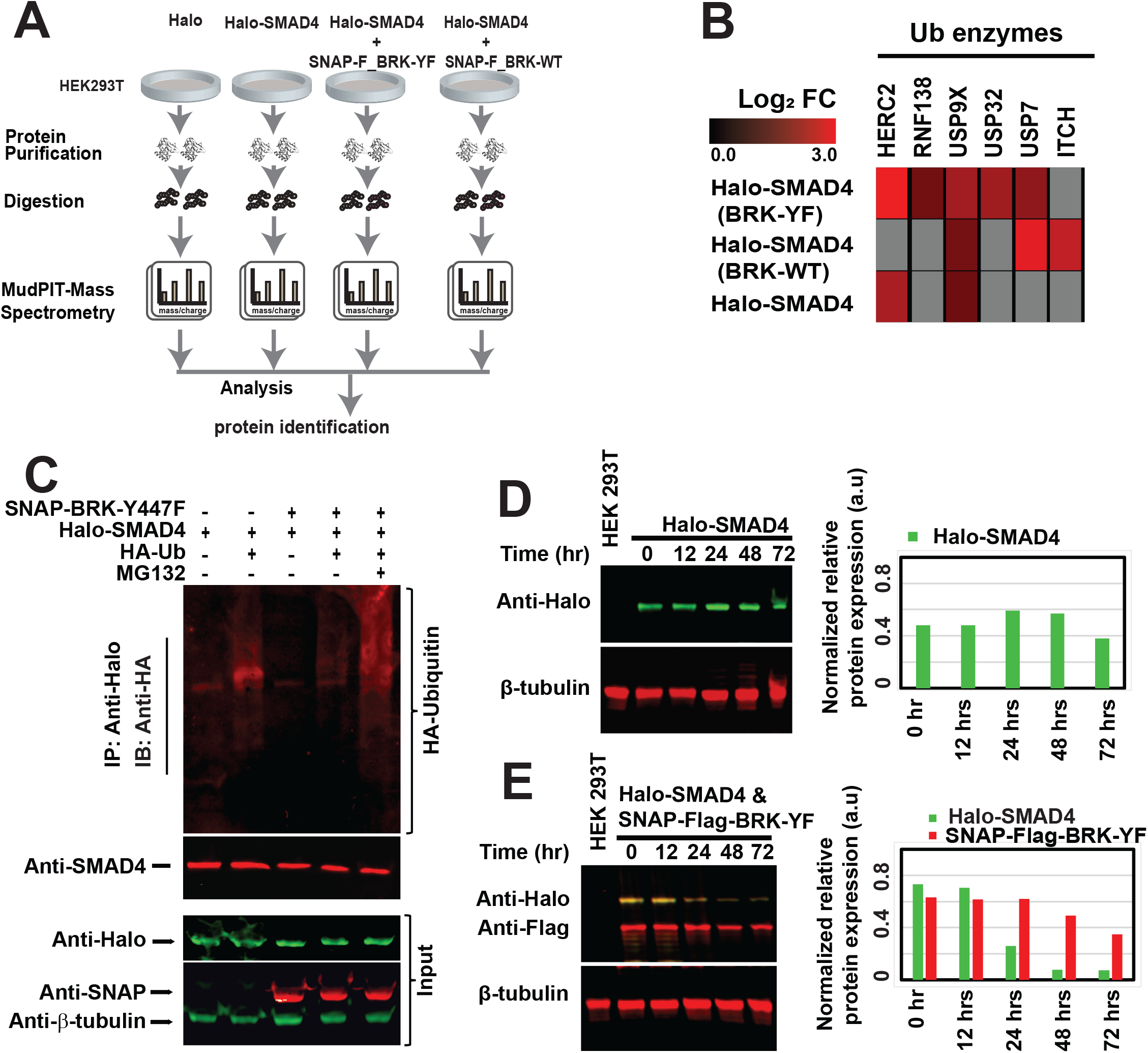
Tyrosine phosphorylated SMAD4 interacts with enzymes of the ubiquitin pathway. **A**. A workflow for discovery proteomics by using MudPIT mass spectrometry for protein identifications. **B. Interaction with ubiquitin modifying enzymes**: relative QSPEC log2 fold changes measured for the specified proteins in the APMS analyses of Halo-SMAD4 with or without BRK-YF were plotted as heat maps (Genesis software package developed by Alexander Sturn and Rene Snajder: http://genome.tugraz.at/genesisclient/genesisclient_description.shtml). **C. SMAD4 polyubiquitination**: HEK293T cells were transiently transfected with Halo-SMAD4, SNAP-Flag-BRK-YF, or in combination with HA-Ubiquitin plasmid. After 36 hours, the cells were treated with 10 µM MG132 for an additional 8 hours. The total cell lysates were subjected to Halo affinity purification followed by immunoblotting with anti-HA and anti-SMAD4 antibodies. **D & E. SMAD4 stability**: Tet-On inducible Halo-SMAD4 plasmid alone or Tet-On inducible Halo-SMAD4 plasmid with SNAP-Flag-BRK-YF were transfected into HEK293T cells for the indicated time points and analyzed by immunoblotting with anti-Halo, anti-Flag, and β-tubulin antibodies. The protein expression was quantified using Image J software and plotted as a bar diagrams.

To investigate the rate of proteasomal degradation of SMAD4 in presence or absence of BRK-YF, we expressed Halo-SMAD4 in HEK 293T cells under the control of a tetracycline-inducible promoter, with or without BRK-YF. After 24 hours post-transfection, tetracycline-containing media was replaced with regular HEK 293T cells culture media and cells were periodically harvested as indicated in **Figure 5D-E**. Immunoblotting of cell lysates showed that Halo-SMAD4 protein levels were dramatically reduced in the presence of activated BRK as early as 24 hours (**Figure 5E**), while in the absence of BRK-YF, Halo-SMAD4 protein levels had not significantly decreased even after 72 hours (**Figure 5D**). This difference in protein stability hence consistent with a role for BRK in regulating SMAD4 degradation.

In summary, our data indicate that ubiquitin ligases are recruited to phosphorylated-SMAD4 in the presence of BRK-YF, leads to its degradation by the proteasome.

### BRK-mediated phosphorylation of Tyrosine 353 and 412 of SMAD4 is required for ubiquitination and degradation of SMAD4

To gain further mechanistic insight, we again used Halo-MudPIT APMS to explore the BRK-YF-modulated SMAD4 ubiquitination and degradation using a mutant of SMAD4 (Halo-SMAD4 Y353F & Y412F) that could not be phosphorylated on tyrosine residues Y353 and Y412 (**Figure 6A**). As we have described previously with Halo-SMAD4, we analyzed the Halo-SMAD4 Y353F & Y412F associated proteins with or without BRK-YF and established that baits were present in all affinity purifications (**Suppl. Fig. S5A-C**). Interestingly, we found that in the presence of the BRK-YF, RNF138 (a ubiquitin ligase) interacted with Halo-SMAD4 but not with SMAD4 Y353F & Y412F, suggesting that phosphorylation of Y353 and/or Y412 is necessary for this interaction. Similarly, SMAD4 interaction with the ubiquitin ligase HERC2 was significantly augmented in the presence of activated BRK, and this interaction was completely lost with mutant Halo-SMAD4 Y353F & Y412F (**Figure 6B and Suppl. Table 2**). Curiously, the ubiquitin ligase ITCH showed a stronger affinity for Halo-SMAD4 Y353F & Y412F in comparison to Halo-SMAD4 in the presence of activated BRK and did not interact in the absence of BRK. Considering the deubiquitinases, we noticed that USP9X and USP7 showed higher affinity for Halo-SMAD4 Y353F & Y412F in the presence of activated BRK and that the deubiquitinase USP32 interacted only with Halo-SMAD4 in the presence of activated BRK (**Figure 6B**). Interestingly, like BRK, USP32 is also often overexpressed in breast cancer cells and tumors, and its inhibition reduces cell proliferation, migration, and apoptosis(28).

**Figure 6.**
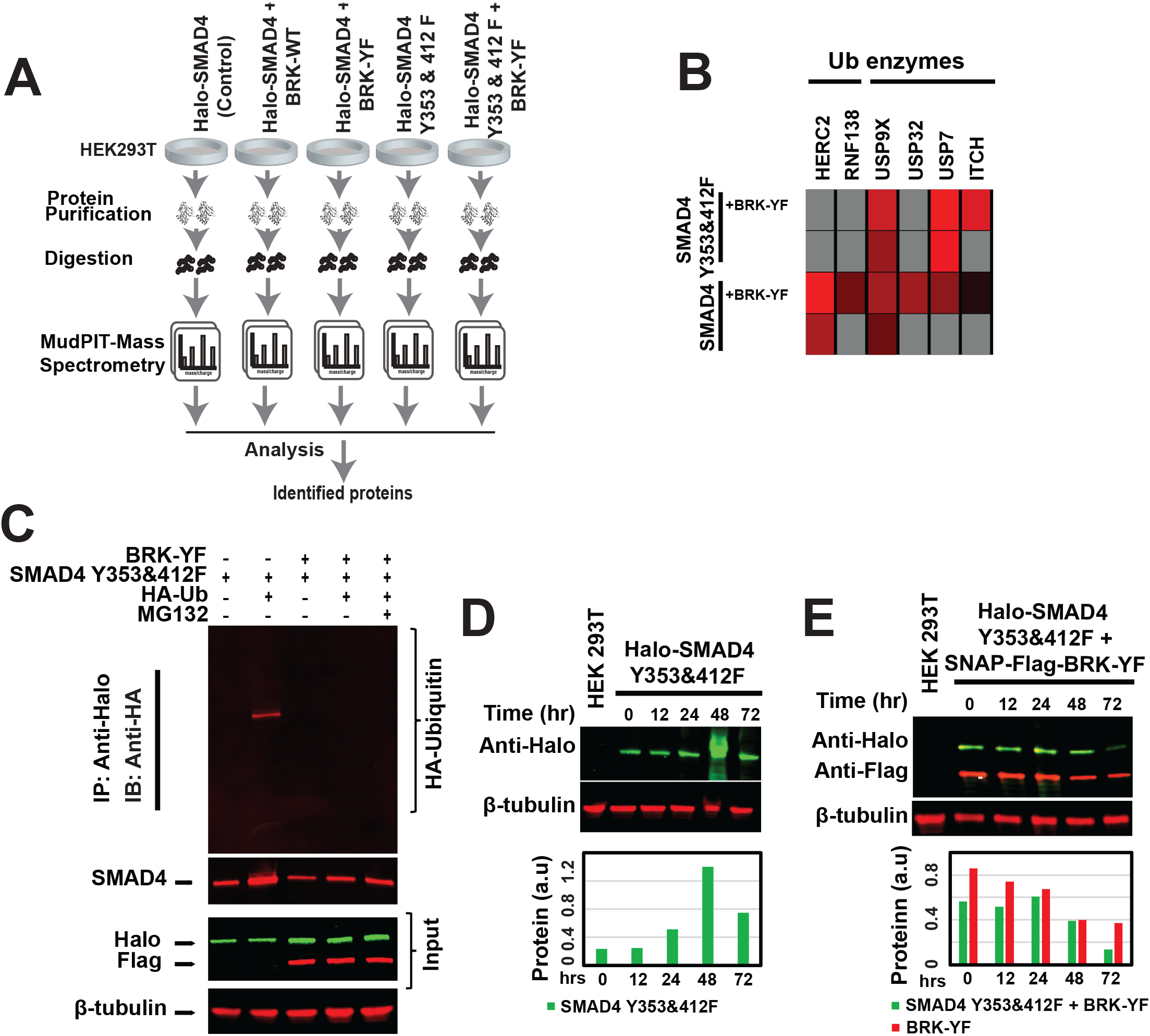
BRK-mediated phosphorylation of SMAD4 is required for its interaction with ubiquitin modifying enzymes. **A**. The experimental flow diagram of discovery proteomics by using MudPIT mass spectrometry for protein identifications. **B. Interaction of Halo-SMAD4 Y353F & Y412F with ubiquitin modifying enzymes**: heat maps of the relative QSPEC-log2 fold changes for the proteins of interest defined in **Figure 5B** measured in the APMS analyses of Halo-SMAD4 or Halo-SMAD4 Y353F & Y412F, presence or absence of BRK-YF. **C. SMAD4 Y353F & Y412F polyubiquitination**: HEK293T cells were transiently transfected with Halo-SMAD4 Y353F & Y412F or SNAP-Flag-BRK-YF or in combination with HA-Ubiquitin plasmid and analyzed as described in **Figure 5C**. **D & E**. **SMAD4 stability**: Tet-On inducible Halo-SMAD4 Y353F & Y412F plasmid with or without SNAP-Flag-BRK-YF were transfected into HEK293T cells and analyzed as in **Figure 5D-E**.

Since our proteomics data (**Figure 6B**) showed that phosphorylation of Y353 & 412F of SMAD4 is essential for ubiquitin ligase recognition, we tested whether the mutant SMAD4 escaped BRK-regulated proteasomal degradation. Plasmids expressing Halo-SMAD4 Y353F & Y412F, SNAP-Flag-BRK-YF, and HA-ubiquitin, were transfected into HEK 293T cells. The transfected cells were treated with MG132 for 8 hours to inhibit 26S proteasome. After Halo affinity purification, we analysed purified samples for ubiquitinated proteins using an anti-HA antibody. Our data showed that ubiquitin was unable to conjugate with mutant Halo-SMAD4 Y353F & Y412F to mark it for proteasomal degradation (**Figure 6C**). Our findings strongly suggest that tyrosine 353 and 412 of SMAD4 are essential for SMAD4 ubiquitination.

Since the tyrosine mutant of SMAD4 was not ubiquitinated, we next examined the stability of SMAD4 Y353F & Y412F in the presence or absence of BRK-YF. To this end, we transfected tetracycline-inducible Halo-SMAD4 Y353F & Y412F alone or with BRK-YF into HEK 293T cells. We found that Halo-SMAD4 Y353F & Y412F protein levels remained stable for a longer time in the absence of activated BRK (**Figure 6D-E**). We also found that the mutant SMAD4 was more stable in the presence of BRK-YF than the WT protein (compare to **Figure 5E**). These findings are consistent with a model whereby BRK-mediated phosphorylation of SMAD4 accelerates its proteasomal degradation, while the phosphorylation-incompetent mutant SMAD4 Y353F & Y412F escapes ubiquitination and subsequent degradation.

### BRK represses the FRK tumor suppressor in a SMAD4-dependent manner to induce EMT and cell invasion

Since SMAD4 is a transcription factor and phosphorylated SMAD4 interacts with chromatin remodeling complexes (Suppl. Fig. 6A&B and Suppl. Table 3), we next investigated genes that might be targeted by SMAD4. We performed an *in-silico* analysis to identify potential SMAD4 binding sites in the genome. We found putative SMAD4-binding sites in the promoter of several genes including tumor suppressor *FRK*(29). Interestingly, three putative SMAD4 binding sites were identified in the *FRK* promoter region (**Figure 7A and Suppl. Fig. 7A**). This finding spurred our interest to further investigate a potential connection between BRK-mediated SMAD4 regulation of FRK expression. To test our hypothesis that SMAD4 regulates this promoter, we performed a luciferase reporter assay and found that luciferase activity was twofold higher in lysates from cells co-expressing SMAD4 and the *FRK* promoter, suggesting that SMAD4 positively regulates the promoter activity of *FRK* (**Figure 7B**), consistent with the presence of SMAD4 binding sites in the *FRK* promoter. Next, we compared the mRNA expression of *FRK* in MDA-MB 231 cells (which express SMAD4, but not BRK – see **Figure 4A**) with *FRK* expression in MDA-MB 231 cells stably expressing BRK-YF. Consistent with BRK-YF-mediated degradation of SMAD4, *FRK* mRNA levels were very low in the cells stably expressing BRK-YF. However, overexpressing SMAD4 in the MDA 231-BRK-YF cells restored expression of *FRK* mRNA (**Figure 7C**). Additionally, we observed that stably expressing BRK-YF decreased the FRK protein levels in MDA 231 cells (**Figure 7D**, compare lanes 1 and 2) and that FRK protein levels were restored by overexpressing SMAD4 (Figure 7D).

**Figure 7.**
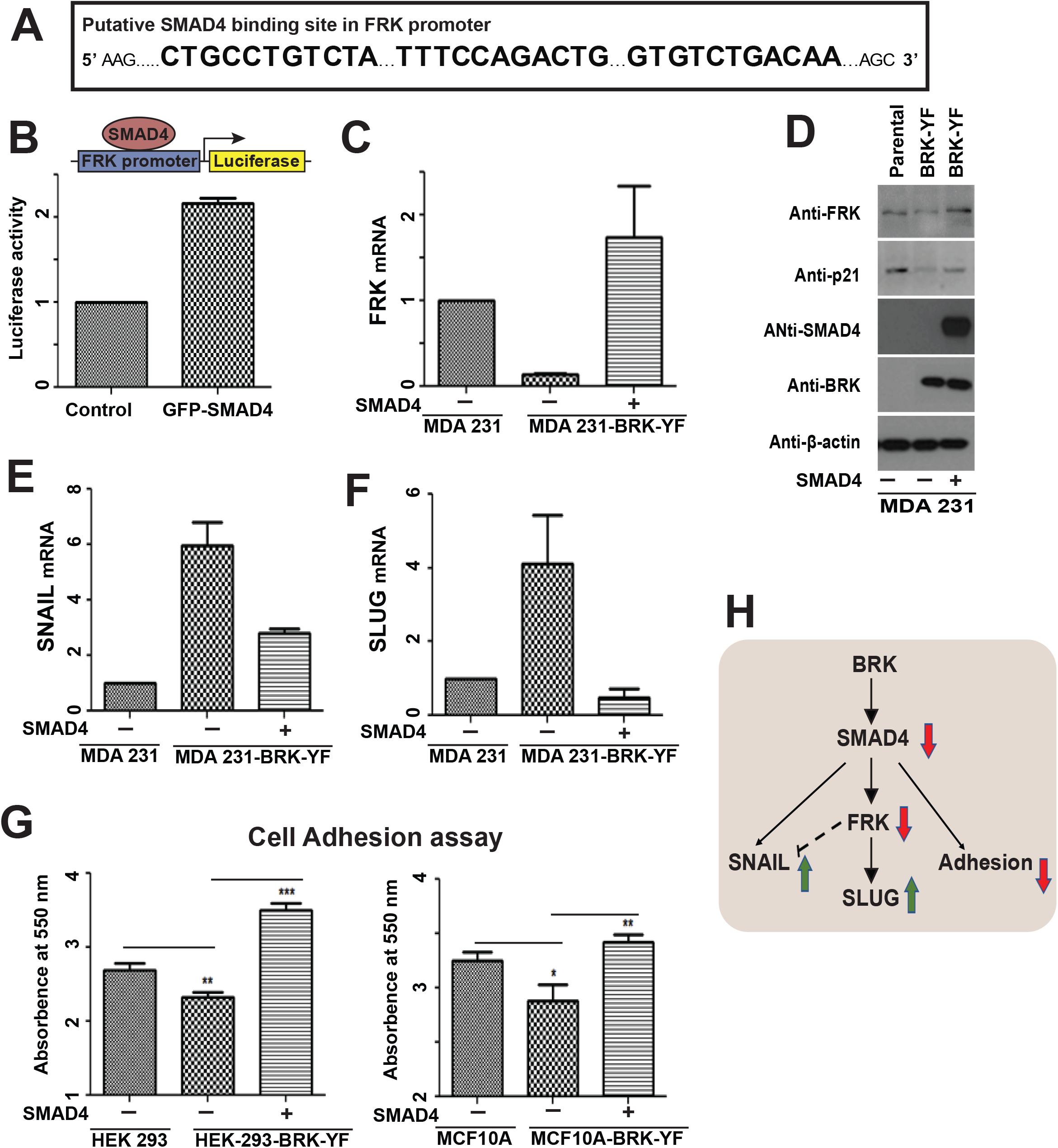
BRK regulates tumor suppressor, EMT markers, and metastatic potential in a SMAD4-dependent manner. **A**. *In silico* analysis shows three putative SMAD4 binding sites in *FRK* promoter. **B**. Luciferase reporter constructs were transfected in HEK293 cells with and without SMAD4 to measure the transcriptional activation of the *FRK* promoter. **C & D**. The mRNA levels of *FRK* were quantified via quantitative RT-PCR and protein levels were analyzed by immunoblotting of the total proteins extracted from parental, stably expressing BRK-YF, and SMAD4-transfected BRK-YF expressing MDA-MB 231 stable cell lines. **E & F**. The mRNA levels *of SNAIL* and *SLUG* were quantified via quantitative RT-PCR in the parental cell line, BRK-YF stable expressing MDA-MB 231 cell line and ectopically expressed SMAD4 in BRK-YF expressing stable cell line. **G**. Cell adhesion assay shows the cell adhesion properties of HEK293 and MCF10A, BRK-YF stably expressing HEK 293 and MCF10A cell lines and SMAD4-transfected stable cell lines. **H**. Activated BRK regulates EMT markers (SNAIL and SLUG) and cell adhesion by modulating SMAD4-FRK.

It has been reported that, FRK suppresses EMT and inhibits cancer metastasis(30) while TGF-β/SMAD4 signaling is crucial for EMT, and promotes metastasis(31). We deduced that the SMAD4-dependent control *of FRK* expression might consequently be controlling the expression of EMT markers. To this end, we examined the expression of EMT markers (E-cadherin, N-cadherin Twist, vimentin, fibronectin, slug, and snail) in parental and BRK-YF expressing MDA-MB 231 cells. Interestingly, we found that *SLUG* and *SNAIL* expression increased in MDA-MB 231 cells stably expressing BRK-YF in comparison to parental cells (six and four folds higher respectively; **Figure 7E-F**). Additionally, SLUG but not SNAIL was significantly suppressed in stably FRK-YF expressing MDA-MB 231 cells (**Suppl. Fig. 7B, C & D**). Moreover, ectopically expressed SMAD4 suppressed the BRK-YF-mediated induction *of SNAIL* and *SLUG* (**Figure 7E-F**). We did not observe any change in the other EMT markers tested (data not shown). Lastly, since our proteomics data showed that the presence of BRK-YF reduced the interaction of SMAD4 with cell adhesion molecules (Suppl. Table 2), we examined the cell adhesion properties of the BRK-YF expressing HEK 293 and MCF 10A cells. Interestingly, we found that activated BRK reduced the cell adhesion capability of BRK-YF expressing cells and that adhesion capability could be completely restored by overexpressing SMAD4 (**Figure 7G**). Overall, our data suggest that activated BRK induces a SMAD4-dependent suppression of tumor suppressor FRK resulting in the stimulation of EMT, and potentially metastasis (**Figure 7H**).

## Discussion

The cellular role of TGF-β/SMAD signaling pathway is a paradox in cancer. On the one hand, *SMAD4*-deficient *Kras^G12D^* pancreatic mouse models showed rapid development of pancreatic tumors(32), while restoration of SMAD4 induced apoptosis(32) and inhibited tumorigenesis in Smad4-defective cancer cells(33). On the other hand, knockdown of SMAD4 significantly reduced liver tumorigenesis in mice(34). The molecular mechanism of this duality is yet to be solved. Further, it was thought that ubiquitination, but not phosphorylation, may play a role in the regulation of SMAD4 function(35). Our current study demonstrates that activated BRK phosphorylates SMAD4 and regulates its stability.

In this study, we have shown that: 1) activated BRK regulates TGF-β/SMAD signaling by interacting with SMAD2/3 and 4; however, SMAD4 is the preferred target of BRK as shown by the results of competitive binding (**Figure 2**); 2) activated BRK phosphorylates SMAD4 on Tyrosines 353 and 412 (**Figure 3**); and phosphorylated SMAD4 is the target of ubiquitin/deubiquitin ligases and is degraded by the ubiquitin-proteasome pathway (**Figures 5-6**); 4) the phosphorylation-regulated degradation of SMAD4 correlates well with the observation that BRK and SMAD4 show inverse expression patterns at the protein levels in breast cancer cells and tumors (**Figure 4**); 5) activated BRK modulates SMAD4 to suppress the tumor suppressor FRK, reduces the interaction of SMAD4 with cell adhesion molecules, and induces EMT (**Figure 7**). Our study provides experimental evidence that the BRK kinase degrades SMAD4 to suppress tumor suppressor FRK and upregulate EMT markers SNAIL and SLUG.

BRK is expressed in most of the cancer types. Although the expression of BRK is ubiquitous in most breast cancer cells and tumors, activated BRK was only detected in the plasma of breast tumors(22). Thus, in this study, we focused on characterizing activated BRK and its role in signal transduction pathways. A kinome array(14) was used to uncover the signal transduction pathways regulated by activated BRK in cancer and normal cells. Ectopically expressed activated BRK regulates the TGF-β/SMAD signaling pathways by interacting with SMAD2/3 and SMAD4. However, SMAD4 outcompetes both SMAD2 and SMAD3 in a binding assay for activated BRK. The BRK modular SH3 domain mediates its protein-protein interactions to regulate signaling(36). Of note, the SMAD4 activation domain (SAD) is proline-rich and contains the Pro-X-X-Pro motif(37), which is the recognition site for SH3 domain(38). However, our domain truncation data indicate that activated BRK interacts with an MH1 domain of SMAD4, which does not contain the SH3 recognition motif, suggesting that the Pro-X-X-Pro motif might be dispensable for SH3-mediated protein-protein interactions.

Dupont *et al*. previously found that a cycle of ubiquitination and de-ubiquitination regulates the function and protein-protein interaction of SMAD4: Ecto/TIF1γ-mediated mono-ubiquitination disassembles SMAD4 from the SMAD complex, while deubiquitination by FAM/USP9x allows SMAD4 to return to SMAD signaling pool(39). Interestingly, our MudPIT-proteomics data revealed that BRK-phosphorylated SMAD4 interacts with several ubiquitin and deubiquitin ligases such as HERC2, RNF138 (ubiquitin ligases) and USP32, USP7, USP9X (deubiquitinases). When phosphorylated by activated BRK, SMAD4 becomes a target of ubiquitin ligases (HERC2 and RNF138), which accelerates its degradation by the ubiquitin-proteasome pathway. Recently, SMAD4 has been shown to phosphorylated and degraded (on Thr277) by GSK3(40). However phosphorylated SMAD4 also becomes a target of deubiquitin ligases (USP32, USP7, USP9X). In fact, deubiquitin ligase USP9X shows a stronger affinity for tyrosine phosphorylated SMAD4 than non-phosphorylated SMAD4. Additionally, our evidence reveals that BRK or other protein tyrosine kinases (since protein tyrosine kinase are functionally redundant(41)) mediates phosphorylation of SMAD4, which is required for SMAD complexes to interact with chromatin remodelers such as SWI/SNF, mediator, HATs, or SIN3/HDAC complexes for gene regulation(42), (43). Of note, in our future study we opt to explore the impact of SMAD4-chromatin remodeler complexes interaction on gene expression in normal and cancer cells.

Complete loss or mutation in SMAD4 has been reported in several cancer types including pancreatic, cholangiocarcinoma, and colorectal cancer(44). Additionally, SMAD4 protein levels decline concurrently with the cumulative malignancy of the tumor cells(45). Our data demonstrate differential expression of SMAD4 and BRK in a panel of breast cancer cell lines. Interestingly, an inverse pattern of SMAD4 and BRK protein levels was noticed in HER2+ (HCC1428 and SKBR3) and TNBC (MDA MB 231, BT20, Hs578T, BT549 and HCC1954) cells, but not ER+ cells (T47D, MCF7, and BT474). However, there are no discrepancies in the expression of *SMAD4* and *BRK* at the mRNA level in those cell lines, indicating a post-translational mechanism of regulation of SMAD4. We also found that the inversely-correlated pattern of expression between SMAD4 and BRK in patients’ breast tumor tissues. In agreement with what is observed in TNBC cells, patients breast tumor samples also show higher levels of the BRK protein in comparison to SMAD4. Furthermore, cells stably expressing activated BRK show a drastic suppression of SMAD4 protein levels. We confirmed that p21, a known downstream target of SMAD4(43), is indeed suppressed in these cell lines expressing activated BRK. However, SMAD4 was not restored or upregulated in MCF7 cells where BRK was knocked-down. This suggests that SMAD4 escapes being targeted for proteasomal degradation, which could be due to a mutation or post-translational modification of SMAD4 or inactivation of BRK in MCF7 cells.

The growth inhibitory signals of TGF-β/Smad4 signaling in early stages of carcinogenesis is well established. Particularly the involvement of SMAD4 in EMT process and in cancer progression in later stages of carcinogenesis in largely unclear(46). Moreover, the function of SMAD4 is mostly contextual. Our data indicate that SMAD4 binds in the promoter region and promote FRK expression. However, in the presence of activated BRK, *FRK* expression is repressed in TNBC cells. Interestingly, overexpression of SMAD4 restores the BRK-induced suppression of FRK level in the TNBC cells stably expressing BRK. Previous studies have shown that the expression of *SNAIL* is upregulated in FRK-knocked down breast cancer cells(30). We observed that activated BRK induces *SLUG* and *SNAIL* expression, while overexpression of FRK significantly suppresses the mesenchymal marker *SLUG*, suggesting an FRK-dependent mechanism for BRK induced promotion of EMT.

In summary, we provide additional evidence to counter the long-standing idea that SMAD4 is not regulated by phosphorylation. We have found that activated BRK competitively binds and phosphorylates SMAD4 and regulates TGF-β/SMAD signaling pathways. Phosphorylated SMAD4 becomes a target of ubiquitin ligases subsequently degraded through the ubiquitin-proteasome system leading the suppression of tumor suppressor FRK. Activated BRK also reduces cell adhesion ability and induces EMT in a SMAD4-dependent manner. Thus, our data suggest that combination therapies targeting activated BRK signaling may have synergized the benefits in the treatment of SMAD4 repressed cancers.

## Material and Methods

### Antibodies and reagents

The following antibodies were obtained from Santa Cruz Biotechnology (Santa Cruz, CA, USA): anti-BRK (sc-916), anti-phosphotyrosine PY20 (Sc-508), anti-SMAD2/3/4, anti-tubulin (Sc-9104), anti-GFP (Sc-8334), and anti-β-actin (sc-130300). Anti-α-Tubulin mouse monoclonal (T9026) antibody was purchased from Sigma-Aldrich (St. Louis, MO). Anti-Halo rabbit polyclonal antibody (G9281) and Magne™ HaloTag^®^ magnetic affinity beads were purchased from Promega (Madison, WI). Proteasome inhibitor MG132 was obtained from Sigma-Aldrich (St. Louis, MO).

### Cell cultures

MCF-10A, MCF12F, MDA-MB-231, BT20, HCC1428, SKBR3, Hs578T, T47D, BT549, MCF7, BT474, HCC1954, and HEK 293 cells were purchased from and cultured according to the American Type Culture Collection (ATCC, Manassas, VA, USA).

### Construction of expression plasmids in human cells

The pGFP-C1-Smad2, -Smad3, and -Smad4 plasmids were a gift from Dr. Caroline Hill, Cancer Research UK. SMAD4 and BRK-YF were subcloned by inserting PCR products (containing SgfI and PmeI restriction sites) to generate pcDNA5-Halo-SMAD4 and pcDNA5-Halo-BRK-YF. Vectors expressing Halo-pcDNA5-Halo-SMAD4_1-140, pcDNA5-Halo-SMAD4_1-320, pcDNA5-Halo-SMAD4_140-320, pcDNA5-Halo-SMAD4_320-552, and pcDNA5-Halo-SMAD4_140-552 were constructed by inserting PCR products between the SgfI and PmeI restriction sites. We further subcloned SMAD4 into pcDNA™5/FRT/TO Vector (a generous gift from the Conaway Lab at Stowers Institute) using Gibson Assembly^®^ Cloning Kit (NEB). Human SMAD4 double mutant (Tyr 353 Phe and Tyr 412 Phe) was obtained by using PCR and Gibson Assembly^®^ Cloning Kit (NEB). All plasmid constructs were confirmed by sequencing. The primers were used for cloning and PCR were listed in the supplemental Table 4.

### Generation of stable cell lines

The construction of cell lines stably expressing GFP-BRK-Y447F has been previously described(5). Amphotropic HEK293-derived Phoenix packaging cells were used to package pBabe-puro retroviral system. For retrovirus production, packaging cells were cultured in DMEM supplemented with 10% bovine calf serum. Transfection with 1% PEI (Polysciences Inc) was conducted with 10 μg of retroviral DNA in 60 μl of 1% PEI plus 430 μl of 0.15M NaCl for the 100 mm culture plates. After 24 h and 48 h, the virus-containing supernatant was collected and filtered through 0.45 μm syringe filter, aliquoted and stored at −80 °C. To infect MCF10A, MDA-MB-231, and HEK293 cells, virus-containing supernatant was supplemented with bovine calf serum and polybrene (Sigma-Aldrich St. Louis, MO) and overlaid on the cells. After overnight incubation, the viral supernatant was replaced with fresh culture medium. Pools of GFP-BRK-Y447F expressing cells were selected with puromycin (Sigma-Aldrich). Expression of GFP-tagged BRK-Y447F was detected after 48-72 h of infection by fluorescence microscopy. To produce stable BRK knockdown cell lines, we used BRK-expressing MCF7 parental cell lines. This knockdown experiment was performed according to the manufacturer’s protocol by using shRNA lentiviral vector plasmids from Santa Cruz Biotechnology. The shRNA plasmids generally consisted of a pool of three to five lentiviral vector plasmids, each encoding target-specific 19–25 nt shRNAs designed to knockdown gene expression. As controls, MCF7 cells were infected with a control shRNA and a GFP-control plasmid for transfection efficiency. Transfected cells were selected using puromycin (Sigma-Aldrich).

### Kinome array

High-throughput kinome assay was performed according to the published protocol(14). In brief, the MDA-MB-231, MCF10A, and HEK 293 cells stably expressing GFP-BRK-YF were cultured to ∼80 % confluency in 10 cm culture plates. The cells were harvested and lysed with 100 μL lysis buffer [20 mM Tris-HCl pH 7.5, 150 mM NaCl, 1 mM EDTA, 1 mM ethylene glycol tetraacetic acid (EGTA), 1% Triton, 2.5 mM sodium pyrophosphate, 1 mM Na3VO4, 1 mM NaF, aprotinin 1 g/ml, leupeptin 1 μg/ml and 1 mM phenylmethylsulphonyl fluoride (PMSF)] and incubated on ice for 10 min followed by centrifugation at maximum speed in a microcentrifuge for 10 min at 4 °C. A 70-μL aliquot of clear cells lysate was mixed with 10 μL of activation mix (50% glycerol, 50 uM ATP, 60 mM MgCl2, 0.05% v/v Brij-35, 0.25 mg/mL BSA, and incubated on the peptide array in a humidity chamber for 2 hours at 37 °C. Arrays were then washed with PBS containing 1 % Triton. Slides were submerged in phosphospecific fluorescent ProQ Diamond Phosphoprotein Stain (Invitrogen) with agitation for 1 h. Arrays were then washed three times in destain containing 20 % acetonitrile (EMD Biosciences, VWR distributor, Mississauga, ON) and 50 mM sodium acetate (Sigma-Aldrich) at pH 4.0 for 10 min. A final wash was done with distilled deionized H2O. Arrays were air-dried for 20 min then centrifuged at 3009g for 2 min to remove any remaining moisture from the array. Arrays were analyzed using a GenePix Profes-sional 4200A microarray scanner (MDS Analytical Technologies, Toronto, ON, Canada) at 532–560 nm with a 580 nm filter to detect dye fluorescence. Images were collected using the GenePix 6.0 software (MDS) and the spot intensity signal collected as the mean of pixel intensity using local feature background intensity background calculation with the default scanner saturation level. Data were processed by using the PIIKA2(47) platform (http://saphire.usask.ca/saphire/piika/).

### Preparation of cell lysates

Confluent or sub-confluent cells were harvested and washed with ice-cold PBS (twice). The whole procedures were carried out at 4°C (on ice) unless specified otherwise. Cells were resuspended in freshly prepared lysis buffer (20 mM Tris ph 7.5, 1% Triton, 150 mm NaCl, protease inhibitors: Aprotinin 5 mg/l and PMSF 0.1 mM) and kept on ice for 30 minutes followed by centrifugation at 14, 000 rpm for 15 minutes at 4°C. Cells were directly lysed in SDS sample buffer [50 mM Tris/HCl (pH 6.8), 2% SDS, 0.1% Bromophenol Blue and 10% glycerol] to obtain total-cell lysates.

### Live cell imaging

HEK 293T cells were seeded onto glass-bottom culture dishes (MatTek, Ashland, MA) and transiently transfected with the SNAP-Flag-BRK and Halo-SMAD4 constructs. Affinity tagged proteins were fluorescently labeled during growth, either with Halo-Tag TMRDirect ligand (Promega) or SNAP-Cell 505-Star (NEB) or with both ligands according to the manufacturer’s instructions. Images were taken with a Zeiss LSM 780 confocal microscope with argon laser excitation at 573-687nm for TMRDirect and 499-526nm for SNAP-Cell 505-Star. To limit photobleaching, exposure time and laser power were adjusted to enhance image quality. An alternating excitation mode was adopted to eliminate cross-talk between color channels. HaloTag™-SMAD4 or SNAP-Tag BRK-YF were ectopically expressed in HEK293T cells and plated at 20% confluency onto glass-bottom MatTek culture dishes (35 mm, No. 2 14mm diameter glass). To label Halo-SMAD4 proteins the HaloTag™ TMRDirect ligand was added in a final concentration of 100 nM and incubated the cells overnight. Additionally, the SNAP-Cell™ 505 ligands was added directly to the cells to label SNAP-Tag BRK-YF in a final concentration of 5 μM and incubated the cells for 1 hour at 37°C in 5% CO2. For co-localization, Halo-SMAD4 and SNAP-Flag-BRK-Y447F constructs were cotransfected into HEK293T cells, and the cells were labeled as indicated above. The cultured media was replaced with OptiMEM to remove background fluorescence prior to imaging. Cells were stained with Hoechst dye to mark nuclei for 30 min prior to imaging.

### Halo Affinity purification of SMAD4 for proteomic analysis

HEK293T cells (1 × 10^7^) were seeded into a 15 cm tissue cultures plates for 24 hours then DNA constructs encoding Halo or SNAP tagged genes of interest were transfected using Lipofectamine LTX (Thermo Fisher Scientific). After 48 hours post-transfection, cells were harvested and washed twice with ice-cold PBS. The ice-cold PBS washed cells were resuspended in 300 μl mammalian cell lysis buffer (Promega) containing 50 mM Tris·HCl (pH 7.5), 150 mM NaCl, 1% Triton^®^ X-100, 0.1% sodium deoxycholate, 0.1 mM benzamidine HCl, 55 μM phenanthroline, 1 mM PMSF, 10 μM bestatin, 5 μM pepstatin A, and 20 μM leupeptin. Next, the cells were raptured by passing through a 26-gauge needle 5-7 times followed by centrifugation at 21,000 × g for 30 min at 4 °C. The resulting 300 μl cell extracts were collected into a new tube and diluted with 700 μl of TBS (50 mM Tris.HCl pH 7.4, 137 mM NaCl, 2.7 mM KCl). To remove insoluble materials, the diluted cell extracts were further centrifuged at 21,000 × *g* for 10 min at 4 °C. Next, 1000 μL of cell extracts were incubated with magnetic beads prepared from 100 μL Magne™ HaloTag^®^ slurry for overnight at 4 °C. Beads were washed four times (750 μL buffer per wash) with wash buffer (50 mM Tris-HCl pH 7.4, 137 mM NaCl, 2.7 mM KCl, and 0.05% Nonidet^®^ P40) before elution. Proteins were eluted by using elution buffer containing 50 mM Tris-HCl pH 8.0, 0.5 mM EDTA, 0.005 mM DTT, and 2 Units AcTEV™ Protease (Thermo Fisher Scientific) for 2 hours at room temperature. The eluate was further passed through a Micro Bio-Spin column (Bio-Rad, Hercules, CA) to remove any residual particles of beads prior to proteomic analysis.

### MudPIT analysis for identification of SMAD4 associated proteins and protein complexes

MudPIT Analysis for protein complexes identification was previously described in detail by Banks *et al.* (26). Briefly, trichloroacetic acid (TCA) precipitated purified proteins were proteolytically digested with endoproteinase Lys-C followed by trypsin digestion, overnight at 37 ° C. A 10-step MudPIT separation approach was applied, and digested peptides were injected directly into a linear ion trap (LTQ) mass spectrometer where spectra were collected and identified. Peptide mass spectra were analyzed by using the ProLuCID(48) and DTASelect(49) algorithms. Next, we used Contrast(49) and NSAF7(50) software to rank the putative affinity purified proteins according to their distributed normalized spectral abundance values (dNSAF)(50). We then used QSPEC(51) to identify enriched proteins in experimental samples compared to control samples. Benjamini and Hochberg statistical method(52) was used to calculate false discovery rates (FDRs) from QSPEC parameters suitable for multiple comparisons. Each of the experiments was repeated at least twice unless otherwise stated.

### Luciferase reporter assays

FRK promoter and SMAD4 plasmids were cotransfected in HEK 293 cells using ViaFect^TM^ (Promega Corporation, Madison, WI) according to the manufacturer’s guidelines. In brief, 125,000 HEK 293 cells were seeded with 500 µl fresh media in each well. The HEK 293 cells were co-transfected with FRK promoter (495 ng; Firefly Luciferase) along with phRL-TK (5 ng; Renilla Luciferase) as an internal control. 48 hours post-transfection, cells were harvested, and luciferase activity was measured by using Dual-Luciferase Assay System with the GloMax^^®^^ 96 Microplate Luminometer (Promega Corporation). To examine the impact of SMAD4 on the *FRK* promoter, cells were co-transfected with GFP-SMAD4 plasmid (250 ng) and *FRK* reporters (245 ng) while control cells were co-transfected with FRK promoter construct and an empty vector (pCDNA3, 245 ng/ well) and 5 ng phRL-TK plasmids as an internal control in each well (Invitrogen Canada).

### RNA isolation, and real-time PCR

Total RNA was isolated from MDA MB-231cells by using RNeasy Plus Mini Kit (Qiagen, Mississauga ON). 1.0 μg of total RNA was used to synthesise cDNA by using Bio-Rad Iscript cDNA Synthesis Kit (Bio-Rad, United States). TaqMan probes Hs00176619_m1, Hs00950344-m1, Hs 00195591_m1 and Hs02758991-g1 were used to quantify the expression of *FRK, SLUG, SNAIL* and *GAPDH* as recommended by the manufacturer (Life Technologies, Burlington, ON, Canada). In brief, 0.6 μL of cDNA, 0.5 μL of probes for each target and housekeeping genes and 5 μL of TaqMan(R) Master Mix were added in each well. dH2O was added in each well to make the volume of 10 μL. Probes for target genes and housekeeping genes were labeled with FAM™ and VIC™ dyes, respectively. The expression of both genes was measured within the same well by using an Applied Biosystems™, Step One Plus qRT-PCR machine (Life Technologies, Burlington, ON, Canada).

### Cell Adhesion assay

96 well plates were coated with either fibronectin or collagen I for an hour at 37°C followed by an hour incubation with 0.5% BSA containing blocking buffer. HEK 293 or MCF10A cells were seeded at a density of 4 × 10^5^ and cells were allowed to attach for 45 minutes at 37°C. The cells were washed three times with 0.1% BSA containing culture media; then the cells were fixed with 4% paraformaldehyde. Next, the cells were stained with crystal violet for 10 minutes. After staining, the cells were washed with distilled water for 10 times. Cells were air dried for an hour and solubilized with 2% SDS on an agitator. The absorbance of each well was measured at 550 nm to quantify the adhesion properties of cells.

### Statistical Analysis

For multiple comparisons (qPCR and adhesion assay), One-way ANOVA followed by a post hoc Newman-Keuls test were used by using GraphPad Prism version 5.04, GraphPad Software, San Diego California USA, www.graphpad.com. The results are presented as the mean ± SD, n≥3 unless otherwise stated. P≤0.05 was considered statistically significant.

## Supporting information

## Acknowledgments

We thank Dr. Stephanie E. Kong for her insightful comments. We thank Dr. Caroline Hill, Cancer Research UK for SMAD 2/3/4 plasmids. This work was supported by the Stowers Institute for Medical Research and the National Institute of General Medical Sciences of the National Institutes of Health under Award Number RO1GM112639 to MPW. The content is solely the responsibility of the authors and does not necessarily represent the official views of the National Institutes of Health. In addition, the work was supported by the 2017 College of Medicine Research Award (CoMRAD) offered by Office of the Vice Dean Research (OVDR) at the University of Saskatchewan, Saskatoon, Canada to KEL, and by the National Science and Engineering Research Council of Canada to SN. Original data underlying this manuscript can be accessed from the Stowers Original Data Repository at http://www.stowers.org/research/publications/LIBPB-1340.

## Author Contributions

SM conceptualized and interpreted all experiments. SM, CAB, AS, LF, SN, KEL, and MPW designed the experiments. SM, CAB, YO, ETB, JMB, AS, CGE performed experiments. SM, CAB, ETB, AS, LF, GH, MS, SN, LF, KEL, and MPW analysed the data and interpreted the findings. KEL and MPW guided the research. SM wrote the manuscript with input from CAB, SN, LF, KEL, and MPW.

## Supplementary Figure legends

**Suppl. Fig. 1.**
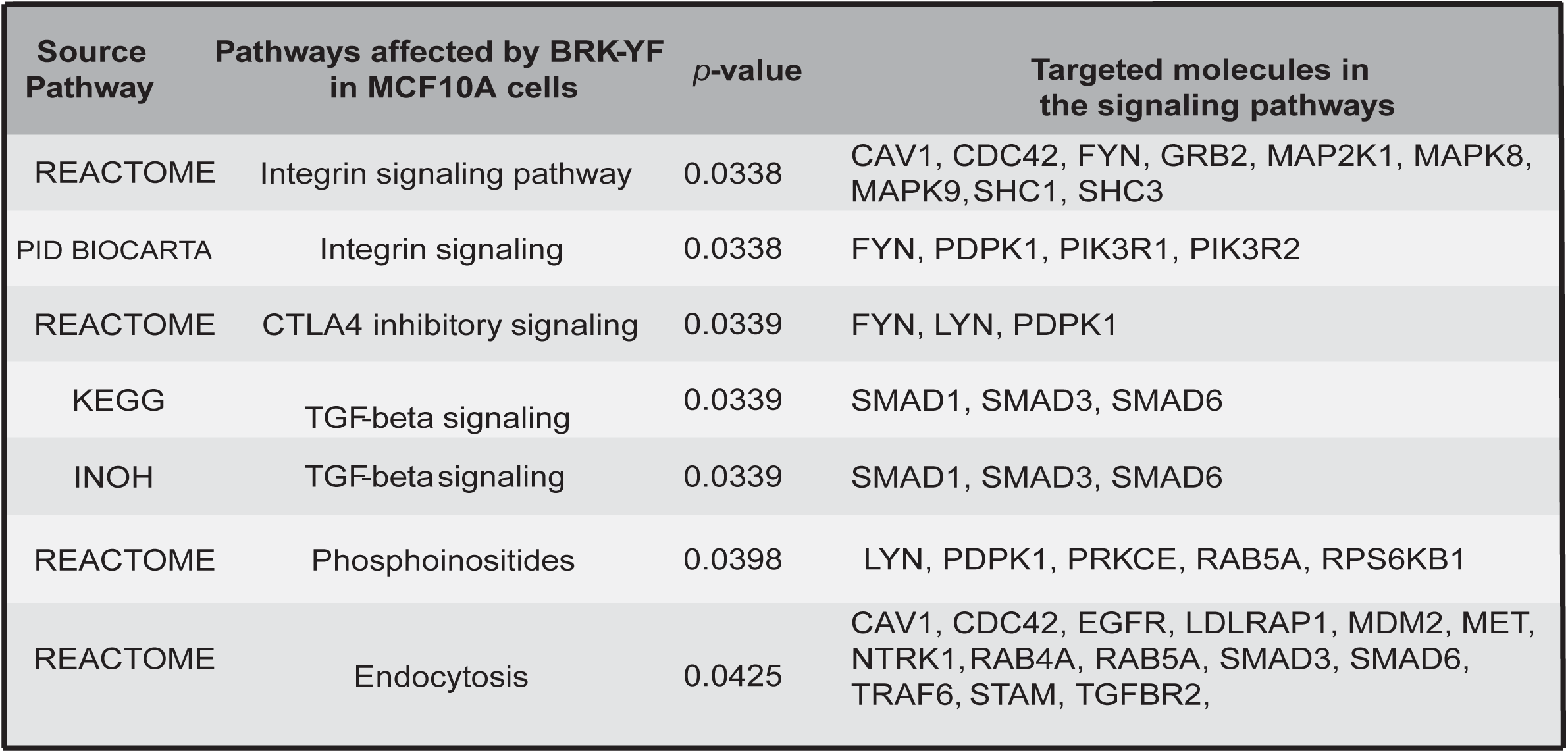
Signaling pathways regulated by activated BRK in MCF10A cells are identified by kinome peptide array analysis (p ≤ 0.05).

**Suppl. Fig. 2.**
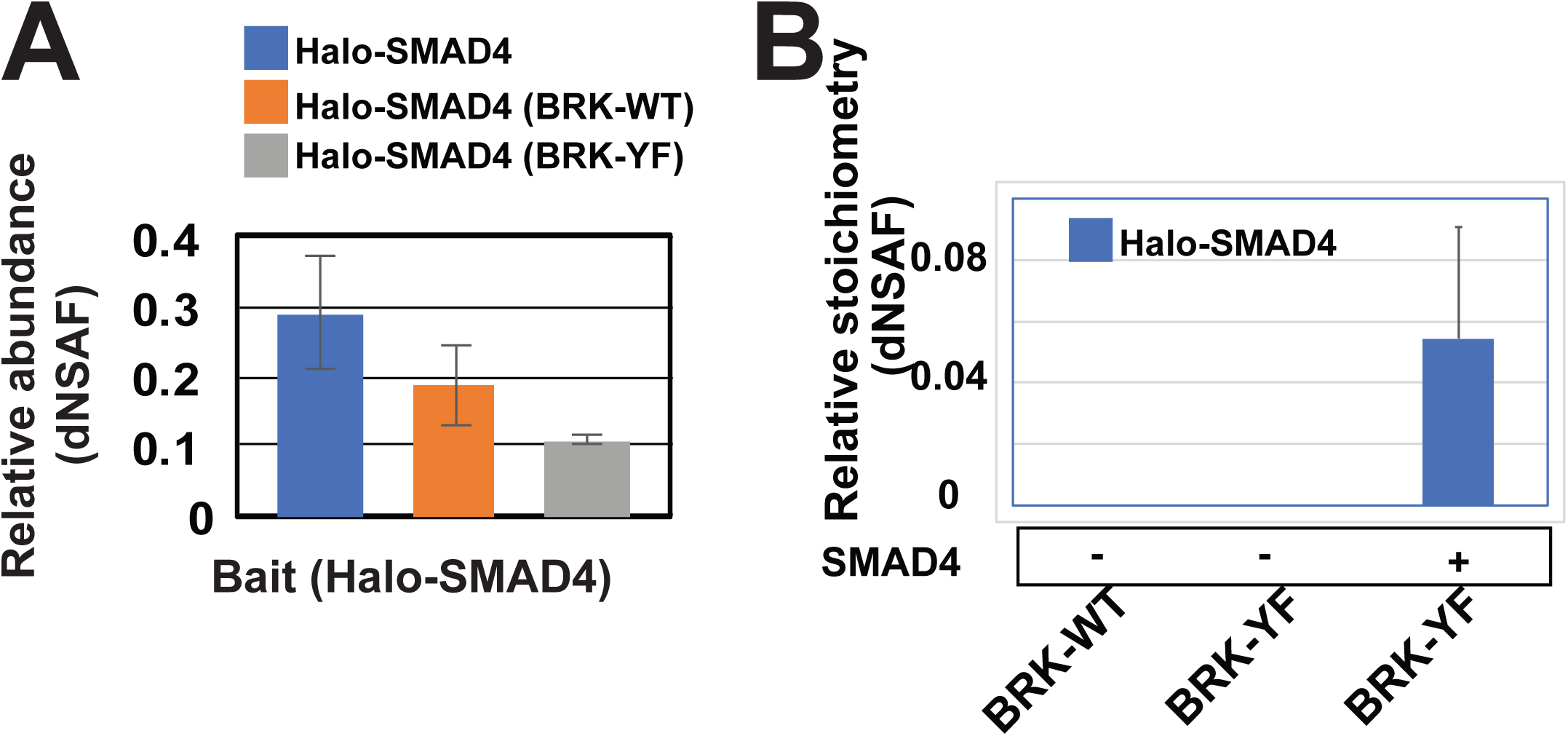
Bait protein abundance. **A**. Bar diagram shows the relative abundance of the bait protein identified by MudPIT analyses of Halo purified Halo-SMAD4. **B**. SNAP-Flag-BRK-WT or SNAP-Flag-BRK-YF with or without Halo-SMAD4 were ectopically expressed in HEK293T cells. Halo affinity purification followed by MudPIT analysis showed Halo-SMAD4 copurified with SNAP-Flag-BRK-YF.

**Suppl. Fig. 3.**
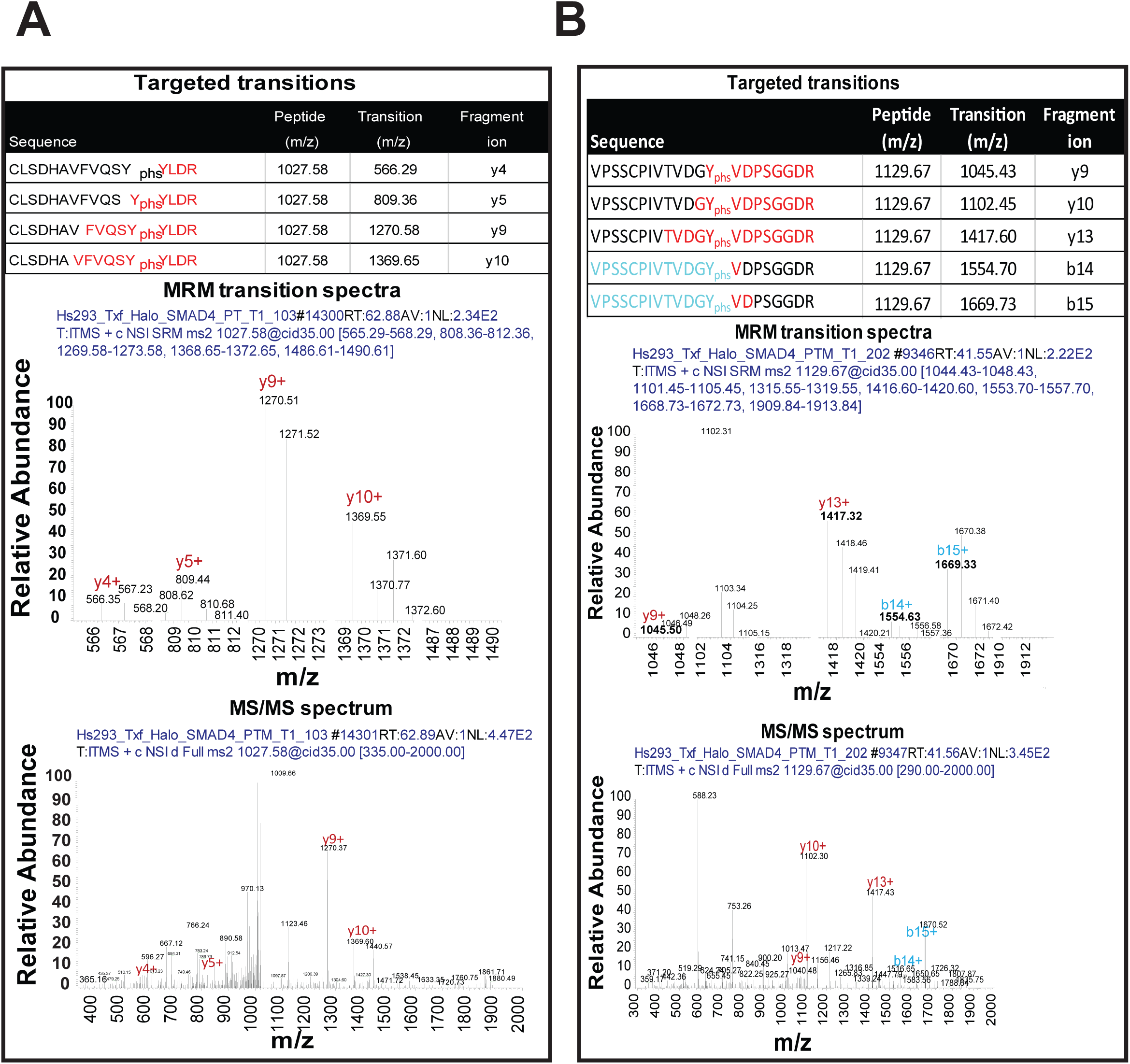
Activated BRK phosphorylates tyrosine 353 and 412 on SMAD4. A & B. Validation of tyrosine phosphorylation on SMAD4 Tyr 353 and 412 in the presence of BRK-YF by multiple reaction monitoring (MRM) approach. The table shows the fragment ions (b and y ions shown in boxes) which were targeted for MRM (Top panel). MRM spectra were acquired when the listed fragment ions were detected (middle panel). The MS/MS spectra were acquired immediately after MRM spectra (bottom panel).

**Suppl. Fig. 4.**
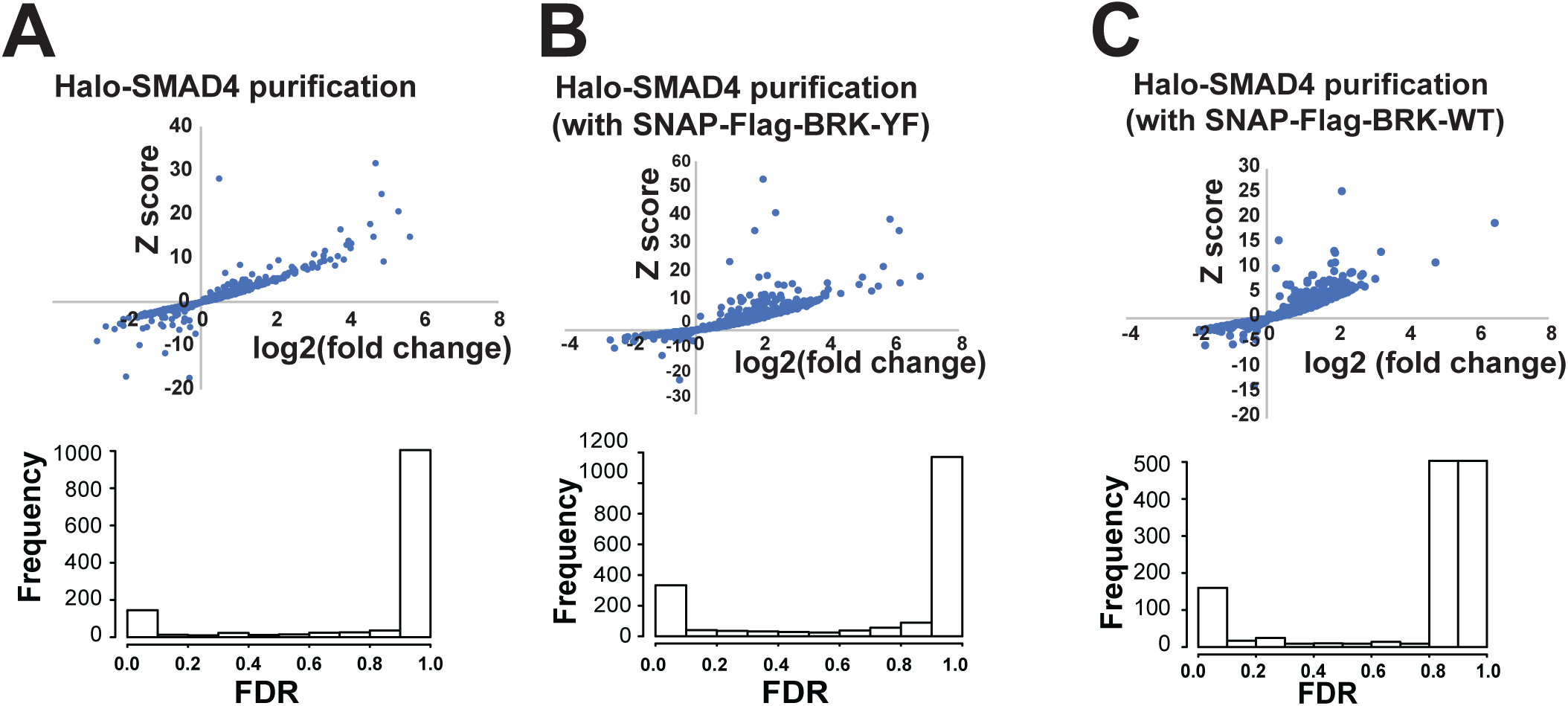
Identification of Halo-SMAD4 associated proteins. A, B & C. Scatterplots show QSPEC log2 FC vs. QSPEC Z score ratios between Halo-SMAD4 and control (Halo alone) experiments. Each dot represents a unique Halo-SMAD4 interacting proteins in presence or absence of activated (BRK-YF) and wildtype BRK (BRK-WT).

**Suppl. Fig. 5.**
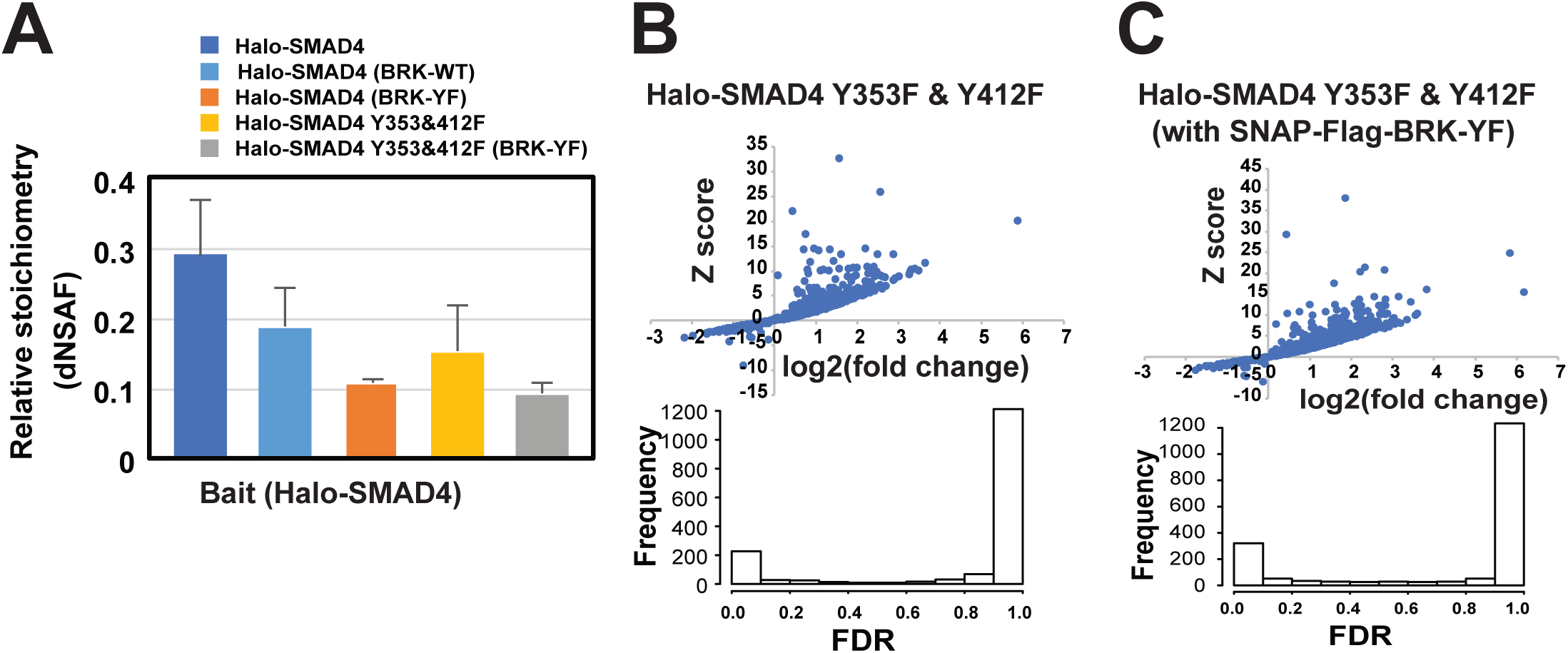
Identification of Halo-SMAD4 Y353F & Y412F associated proteins in presence or absence of BRK-YF. A. Bar diagram showing the relative abundance of bait proteins (Halo-SMAd4 and Halo-SMAD4 Y353F & Y412F) identified by APMS from five different cell lines. **B & C**. Scatterplot is showing the proteins interacting with Halo-SMAD4 Y353F & Y412F in presence or absence of activated BRK.

**Suppl. Fig. 6.**
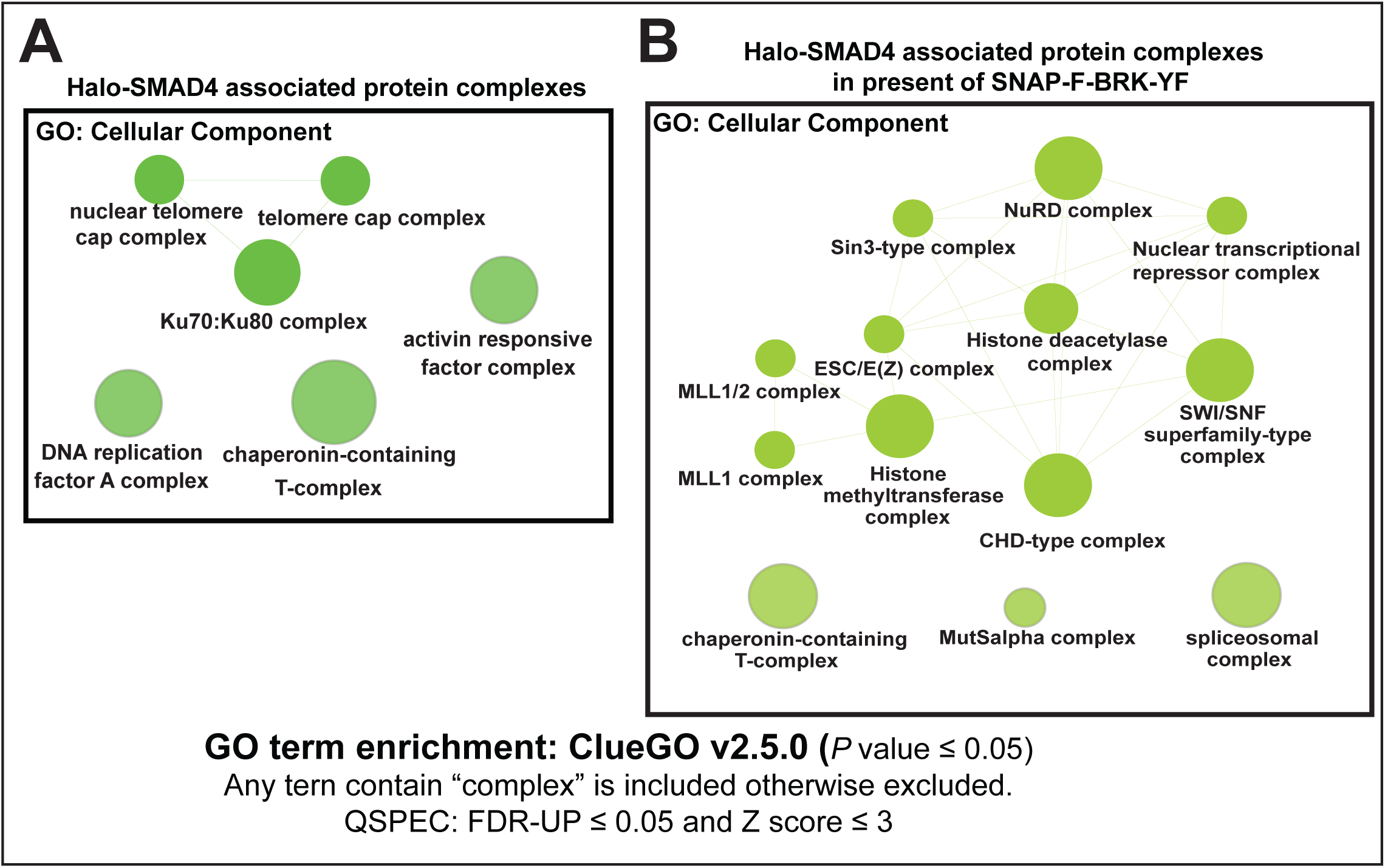
GO analyses for cellular components represented in the proteins associated with Halo-SMAD4 and phosphorylated Halo-SMAD4. Halo-SMAD4 and phosphorylated Halo-SMAD4 associated proteins (Z score ≥ 3; QSPEC FDR ≤ 0.05) were analyzed for a cellular component by using the ClueGo(53) plugin in Cytoscape(54). *P* values ≤ 0.05.

**Suppl. Fig. 7.**
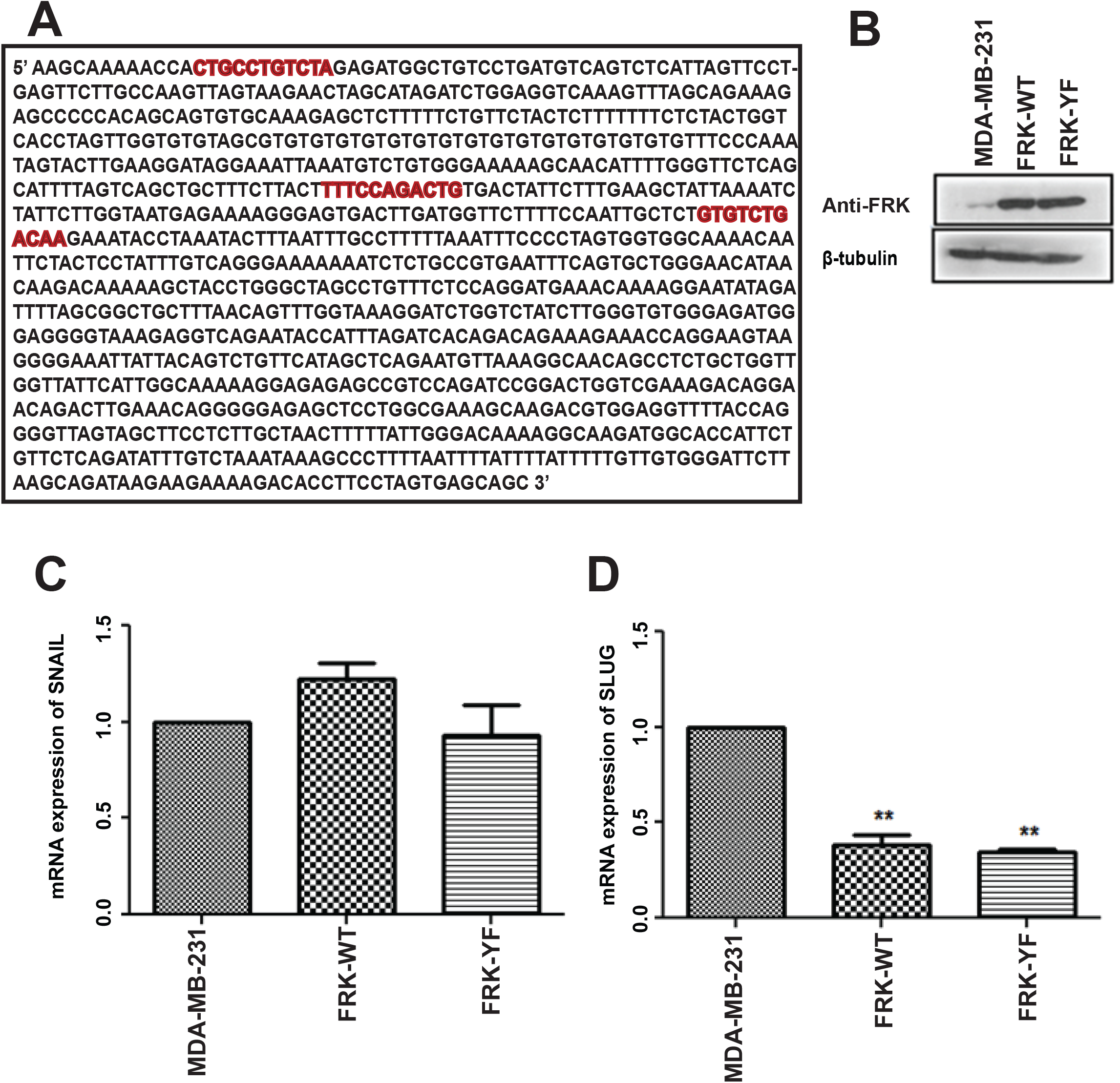
FRK-dependent regulation of EMT markers. A. FRK-WT and FRK-Y497F stably expressing MDA-MB 231 cells were generated, and the expression of FRK-WT and FRK-Y497F was evaluated by immunoblotting. **B & C**. *SNAIL* and *SLUG* mRNA levels were quantified via quantitative RT-PCR in parental, FRK-WT, and FRK-Y497F stably expressing MDA-MB 231 cells.

### Tables

**Suppl. Table 1**. Phosphorylation sites on SAMD4 detected by MudPIT analyses of in presence or absence of BRK-YF.

**Suppl. Table 2**. Differential protein interaction of Halo-SMAD4 in the presence of SNAP-F-BRK-WT or SNAP-F-BRK-YF (QSPEC log2 fold change>= 1 and QSPEC FDR ≤ 0.05). And differential protein interactions of Halo-SMAD4 and Halo-SMAD4 Y353F & Y412F in the presence of SNAP-F-BRK-WT or SNAP-F-BRK-YF (QSPEC log2 fold change>= 2 and QSPEC FDR ≤ 0.05).

**Suppl. Table 3**. GO analysis for cellular component of Halo-SMAD4 in the presence of SNAP-F-BRK-WT or SNAP-F-BRK-YF in ClueGO FDR_0.05; Zscore_3.

**Suppl. Table 4**. Primers

