## Supplementary material for "BRK Phosphorylates SMAD4 for proteasomal degradation and inhibits tumor suppressor FRK to control SNAIL, SLUG and metastatic potential"

**Table S1:** **Phosphorylation sites on SAMD4 detected by MudPIT analyses of Halo-SMAD4 and Halo-SMAD4+BRK-YF**

| | | Halo-SMAD4 | | | | Halo-SMAD4_1 | | | Halo-SMAD4_2 | | | Halo-SMAD4_3 | | |
|---|---|---|---|---|---|---|---|---|---|---|---|---|---|---|
| **Phosphorylated SMAD4 Residue** | **Previously Reported[a]** | **Detected out of 3 Replicates** | **Total_SpC** | **Modified_SpC** | **Modified_SpC: Total_SpC** | **Total_SpC** | **Modified_SpC** | **Modified_SpC: Total_SpC** | **Total_SpC** | **Modified_SpC** | **Modified_SpC: Total_SpC** | **Total_SpC** | **Modified_SpC** | **Modified_SpC: Total_SpC** |
| S138 | x | 3 | 777 | 63 | 8.11 | 234 | 39 | 16.67 | 371 | 8 | 2.16 | 172 | 16 | 9.3 |
| S403 | | 1 | 2957 | 14 | 0.47 | 868 | 0 | 0 | 1126 | 0 | 0 | 963 | 14 | 1.45 |
| S357 | | 2 | 763 | 4 | 0.52 | 222 | 2 | 0.9 | 372 | 0 | 0 | 169 | 2 | 1.18 |
| S517 | | 2 | 226 | 3 | 1.33 | 52 | 2 | 3.85 | 106 | 0 | 0 | 68 | 1 | 1.47 |
| Y513 | x | 2 | 190 | 3 | 1.58 | 51 | 2 | 3.92 | 89 | 0 | 0 | 50 | 1 | 2 |
| S343 | x | 0 | 763 | 0 | 0 | 222 | 0 | 0 | 372 | 0 | 0 | 169 | 0 | 0 |
| S344 | | 0 | 763 | 0 | 0 | 222 | 0 | 0 | 372 | 0 | 0 | 169 | 0 | 0 |
| **Y353** | | 0 | 763 | 0 | 0 | 222 | 0 | 0 | 372 | 0 | 0 | 169 | 0 | 0 |
| **Y412** | | 0 | 2957 | 0 | 0 | 868 | 0 | 0 | 1126 | 0 | 0 | 963 | 0 | 0 |

[a]www.phosphosite.org

| | | Halo-SMAD4_BRK-YF | | | | Halo-SMAD4_BRK-YF_1 | | | Halo-SMAD4_BRK-YF_2 | | |
|---|---|---|---|---|---|---|---|---|---|---|---|
| **Phosphorylated SMAD4 Residue** | **Previously Reported[a]** | **Detected out of 2 Replicates** | **Total_SpC** | **Modified_SpC** | **Modified_SpC: Total_SpC** | **Total_SpC** | **Modified_SpC** | **Modified_SpC: Total_SpC** | **Total_SpC** | **Modified_SpC** | **Modified_SpC: Total_SpC** |
| S138 | x | 2 | 730 | 35 | 4.79 | 520 | 1 | 0.19 | 210 | 34 | 16.19 |
| S403 | | 2 | 1287 | 13 | 1 | 634 | 7 | 1.1 | 653 | 6 | 0.92 |
| S357 | | 0 | 361 | 0 | 0 | 187 | 0 | 0 | 174 | 0 | 0 |
| S517 | | 0 | 24 | 0 | 0 | 11 | 0 | 0 | 13 | 0 | 0 |
| Y513 | x | 0 | 21 | 0 | 0 | 13 | 0 | 0 | 8 | 0 | 0 |
| S343 | x | 1 | 361 | 2 | 1 | 187 | 0 | 0 | 174 | 2 | 1.15 |
| S344 | | 1 | 361 | 2 | 1 | 187 | 0 | 0 | 174 | 2 | 1.15 |
| **Y353** | | 2 | 361 | 3 | 1 | 187 | 1 | 0.53 | 174 | 2 | 1.15 |
| **Y412** | | 1 | 1287 | 1 | 0 | 634 | 1 | 0.16 | 653 | 0 | 0 |

[a]www.phosphosite.org
